# Structures of the active HER2/HER3 receptor complex reveal dynamics at the dimerization interface induced by binding of a single ligand

**DOI:** 10.1101/2021.05.03.442258

**Authors:** Devan Diwanji, Raphael Trenker, Tarjani M. Thaker, Feng Wang, David A. Agard, Kliment A. Verba, Natalia Jura

## Abstract

The Human Epidermal Growth Factor Receptor 2 (HER2) and HER3 form a potent pro-oncogenic heterocomplex upon binding of growth factor neuregulin-1β (NRG1β)^1–3^. The mechanism by which HER2 and HER3 interact remains unknown in the absence of any structures of the complex. We isolated the NRG1β-bound near full-length HER2/HER3 dimer and obtained a 2.9Å cryo-electron microscopy (cryo-EM) reconstruction of the extracellular domain module which reveals unexpected dynamics at the HER2/HER3 dimerization interface. We show that the dimerization arm of NRG1β-bound HER3 is unresolved likely because the apo HER2 monomer fails to undergo a ligand-induced conformational change needed to establish a HER3 dimerization arm binding pocket. In a second structure of an oncogenic extracellular domain mutant of HER2, S310F, we observe a compensatory interaction with the HER3 dimerization arm that stabilizes the dimerization interface. We show that both HER2/HER3 and HER2-S310F/HER3 retain the capacity to bind to the HER2-directed therapeutic antibody, trastuzumab, but the mutant complex does not bind to pertuzumab. Our 3.5Å structure of the HER2-S310F/HER3/NRG1β/trastuzumab Fragment antigen binding (Fab) complex shows that the receptor dimer undergoes a conformational change to accommodate trastuzumab. Thus, like oncogenic mutations, therapeutics exploit the intrinsic dynamics of the HER2/HER3 heterodimer. The unique features of a singly liganded HER2/HER3 heterodimer underscore the allosteric sensing of the ligand occupancy by the dimerization interface and explain why extracellular domains of HER2 do not homo-associate via canonical active dimer interface.

## Introduction

HER2 and HER3 are members of the HER/ErbB family of receptor tyrosine kinases (in addition to EGFR and HER4) that convert the binding of extracellular growth factor ligands into the activation of the intracellular kinase domains. Point mutations and HER2 overexpression have been firmly established as oncogenic in breast, lung, bladder and other tissues, and mutations in HER3 have been identified in colon and gastric cancers^4–11^. HER3 upregulation is also a major mechanism underlying resistance to anti-HER2 treatments^12,13^. To form an active complex, HER receptors dimerize upon binding to growth factor ligands. HER2 is uniquely unable to bind to known growth factors and thus dependent on heterodimerization with other HER receptors for activation and signaling. The preferred dimerization partner of HER2 is HER3, which binds growth factors from the neuregulin (NRG) family^1–3^. Like HER2, HER3 is an obligate heterodimer partner, because it has a catalytically impaired kinase domain (a pseudokinase) and cannot support its own phosphorylation^14,15^.

In the absence of any high-resolution structures of the HER2/HER3 heterocomplex, our current molecular understanding of its activation is mostly inferred from structural studies of the related receptors, EGFR and HER4. On the intracellular side, all HER receptors are composed of short juxtamembrane segments, kinase domains and long unstructured tails. Upon activation, the pseudokinase domain of HER3 is predicted to allosterically activate the HER2 kinase to initiate downstream signaling^16,17^ (**Fig. 1a**). On the extracellular side, HER receptors are composed of four domains (I-IV). Domain II contains the dimerization arm, a key structural element at the dimerization interface (**Fig. 1a**). In the absence of ligand, the dimerization arm is obscured in the inactive “tethered” conformation, as seen in structures of EGFR, HER3 and HER4, by an intramolecular interaction between domains II and IV^18–20^. A critical function of ligand binding is to break the tether and stabilize an extended conformation that exposes the dimerization arm (**Fig. 1a**). While no structures of ligand-bound HER3 have been solved, EGFR and HER4 have been observed in the ligand-bound extended conformation with an exposed dimerization arm which contacts a pocket formed between domains I and III of the other monomer^21–23^. The resulting active extracellular domain dimers of EGFR and HER4 are largely symmetric, stabilized by the binding of two growth factor molecules, with the dimerization arm of each monomer providing significant interaction surfaces^21–24^.

**Fig. 1.**
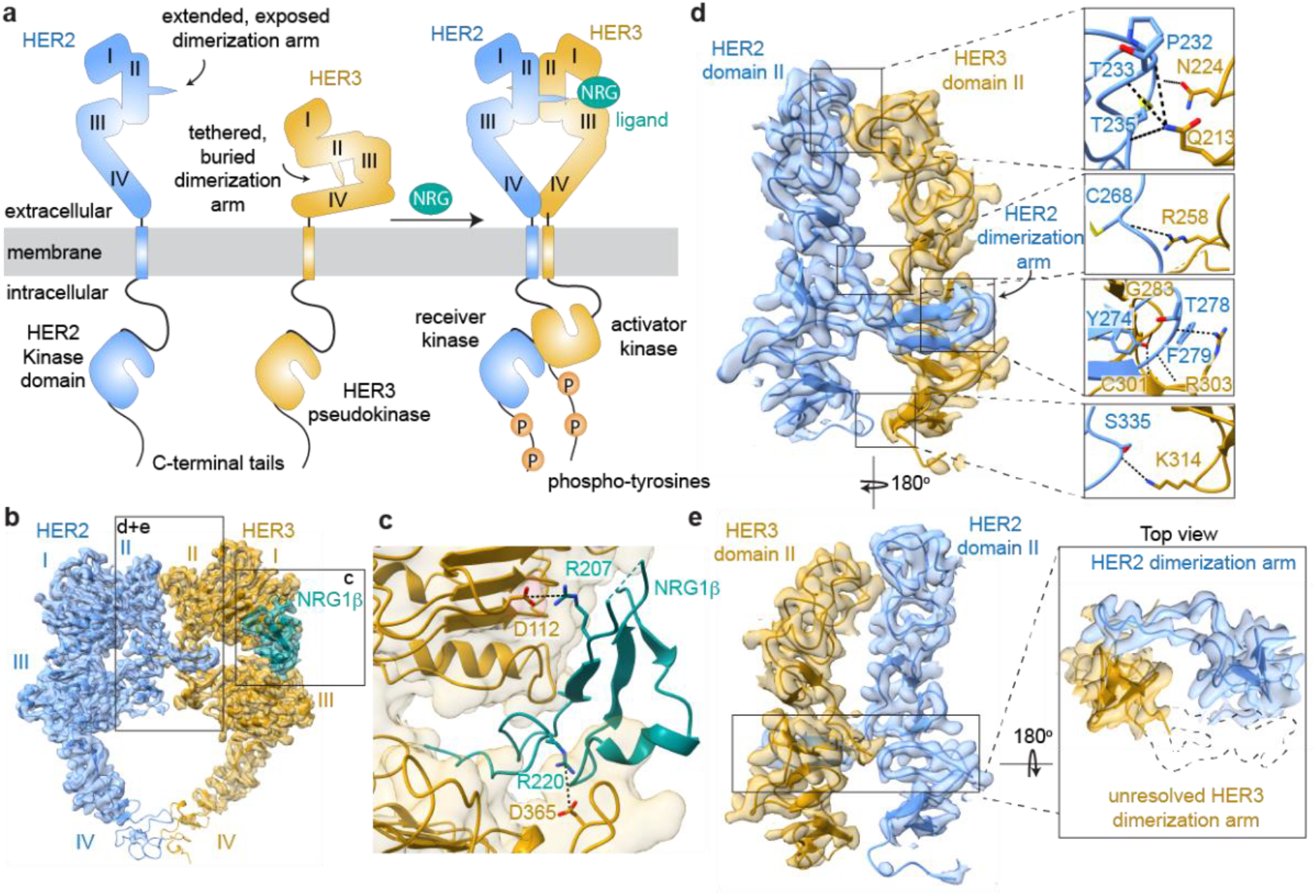
Overall structure of the HER2/HER3/NRG1β extracellular domain complex. **a,** Cartoon schematic of the conformational changes that the inactive HER2 and HER3 monomers are predicted to undergo during heterodimerization in the presence of neuregulin-1β (NRG). **b,** Cryo-EM map and the resulting structural model of the HER2/HER3/NRG1β extracellular domain complex, with HER2 shown in light blue, HER3 in gold and NRG1β in teal. Extracellular domains I-IV are marked on the structures. Boxes indicate insets magnified in c-e. **c,** Detailed view of the NRG1β binding site on HER3. HER3 is shown in cartoon representation and molecular surface, in gold. NRG1β is in teal. Salt bridges are shown in dotted black lines. **d,** Detailed view of the dimerization interface between domains II of HER2 and HER3 with all polar contacts between receptors highlighted in the boxes on the right. **e,** Same view as in d but rotated 180°to illustrate lack of density for the HER3 dimerization arm. An outline of the expected location of the HER3 dimerization arm based on previous extracellular domain structures is shown as a dotted path in the top view.

The orphan receptor HER2 is found in an extended conformation in all current structures despite lacking bound growth factor ligands^25,26^. In contrast to EGFR and HER4, however, homodimeric interactions mediated by the extended extracellular domain of HER2 have not been observed even though its dimerization arm is constitutively exposed. It has been proposed that the existing extended HER2 extracellular domain structures are in a constitutively autoinhibited conformation due to their similarities with the inactive structures of EGFR^27^. Whether HER2 adopts a different, “active” conformation when it binds other HER receptors, and what this conformation may look like, remains a mystery. The stabilization of such HER2-containing complexes by the binding of only one growth factor stands in contrast to all known high-resolution structures of the active EGFR and HER4 receptor extracellular domain homodimers. In this study, we used cryo-EM to gain structural insights into the formation of the active HER2/HER3 complex, the activation mechanism of extracellular domain cancer mutations, and the binding of the heterocomplex to existing HER2-directed therapeutics.

## Results

### Intracellular domains stabilize the ligand-induced extracellular domain interactions in the HER2/HER3 complex but are structurally labile

Previous biophysical studies on the isolated receptor extracellular domain fragments of HER2 and HER3 did not yield stable heterodimeric complexes in the presence of NRG ligands^28^. We hypothesized that the transmembrane and intracellular kinase domains might contribute to the stabilization of extracellular domain interactions, allowing us to reconstitute the HER2/HER3 heterodimeric complex. We expressed near-full length HER2 receptors in the presence of the covalent kinase inhibitor canertinib^29^ and HER3 receptors with only their unstructured intracellular C-terminal tails truncated (see **Methods**, **Extended Data Fig. 1**). After extensive optimization, we were able to obtain the NRG1β-bound near full-length complexes of HER2/HER3 solubilized in detergent in sufficient amounts and homogeneity for cryo-EM. A critical factor for reaching the necessary yields was the inclusion of cancer-associated mutations in the HER3 pseudokinase domain, which we have previously shown increase HER3 dimerization affinity with EGFR^17^. Another major enabling technology was the use of graphene oxide coated holey carbon grids that enabled solving our high-resolution cryo-EM structures at low receptor concentrations^30,31^. While the transmembrane and intracellular kinase domains are essential for the stabilization of the extracellular domain interactions between HER2 and HER3, they do not appear to be rigidly connected to the extracellular domains, and their inclusion in the cryo-EM reconstructions of all three states ultimately limited the resolution of the full-length receptor heterodimer structures (**Extended Data Fig. 1**). However, focusing on the extracellular domain yielded a 2.9Å structure of the HER2/HER3/NRG1β extracellular domain complex (**Fig. 1b**, **Extended Data Fig. 2**).

### Structure of the asymmetric HER2/HER3 dimer reveals that the dimerization arm of HER3 is disengaged

In the cryo-EM reconstruction of the HER2/HER3/NRG1β extracellular domain module, HER2 and HER3 assemble in a “heart-shaped” dimer, resembling a conformation similar to that of the known ligand-stabilized EGFR and HER4 extracellular domain homodimers^21–24,32^ (**Extended Data Fig. 3** and **Extended Data Fig. 4**). Active EGFR extracellular dimers have been observed in highly symmetric complexes when bound to high affinity ligands such has EGF and in slightly asymmetric complexes when bound to lower affinity ligands such as epiregulin (EREG)^21–24,32^. The HER2/HER3/NRG1β complex represents a conformationally distinct HER receptor dimer with the highest degree of asymmetry (**Extended Data Fig. 3** and **Extended Data Fig. 4**). Membrane-proximal domains IV are visualized at lower resolution for both receptors (Extended Data Fig. 2) indicating their flexibility, as observed in EGFR and HER4^22–24,33^. Our structure features the previously uncharacterized extended state of the HER3 extracellular domain, with NRG1β clearly resolved in the density (**Fig. 1b,c**). NRG1β engages with HER3 primarily through an extensive interaction network at its C-terminus (total buried surface area: 2,666 Å^2^) stabilized by salt bridges between R207 (NRG1β) and D112 in HER3 domain I, and R220 (NRG1β) and D365 in HER3 domain III, bringing domains I and III into close proximity (**Fig. 1c**). Many of the contacts between NRG1β and HER3 are conserved in the NRG1β-bound HER4 complex (**Extended Data Fig. 3**).

Like HER3, HER2 adopts an extended conformation in the dimer. This conformation is almost identical to the one previously seen in structures of monomeric HER2 (**Extended Data Fig. 3**) (RMSD: 1.01 Å (1N8Z), 0.74 Å (1N8Y), 0.97 Å (6OGE))^25,26^. As previously noted, this conformation differs from the extended conformations of other HER receptors especially in the curvature of domain II, the presentation of the dimerization arm and the relative orientation of domains I and III. Our structure provides evidence that this atypical extended state of HER2 is readily accommodated in the active HER2/HER3 dimer, contradicting the hypothesis that HER2 needs to undergo additional conformational changes in order to engage with other HER receptors in active dimers^27^.

The most striking feature of the HER2/HER3/NRG1β complex is the lack of resolvable density for the HER3 dimerization arm in domain II (**Fig. 1e**). This is surprising because in all other structures of active HER receptor extracellular domain dimers, the dimerization arms of both receptors provide essential contributions to the dimerization interface. In contrast, only the HER2 dimerization arm is resolved in the HER2/HER3/NRG1β structure and interacts with HER3 through a number of key sidechain-backbone and backbone-backbone interactions between HER2-Y274/HER3-G283, HER2-Y274/HER3-R301, HER2-T278/HER3-R303 and HER2-F279/HER3-R303 (**Fig. 1d**). Apart from the dimerization arm-mediated polar interactions, domain II of HER2 interacts with domain II of HER3 via six additional hydrogen bonds, which while fewer, are similarly positioned to those seen in extracellular crystal structures of other HER receptors (**Fig. 1d**, **Extended Data Fig. 4**). Consequently, the total buried surface area at the HER2/HER3 heterodimer interface (1,821 Å^2^, domains I-III only) is reduced compared to that of other HER homodimers (2,773 Å^2^ for EGFR homodimer bound to EGF, 2,135 Å^2^ for EGFR homodimer bound to EREG and 2,673 Å^2^ for HER4 homodimer bound to NRG1β)^23,24^.

### Ligand binding mode determines the extent of dimerization arm engagement

One possible explanation for why we do not observe the dimerization arm of HER3 in our structure is that HER2 does not have a suitable binding pocket to engage the HER3 dimerization arm. In the symmetric, EGF-bound EGFR extracellular domain dimers each protomer cradles the dimerization arm of the other protomer via an enclosed binding pocket. In comparison, domain I/III interface in HER2 does not fully close, which compromises the binding pocket for the HER3 dimerization arm (**Fig. 2a**). We investigated whether such a weakened, open pocket is a unique feature of HER2 by analyzing known crystal structures of other HER receptor ectodomains (**Fig. 2a**). We noticed that in the asymmetric structure of the EREG-bound EGFR ectodomain dimer, the dimerization arm-binding pocket was also open, although only partially, and only in one receptor monomer^32^ (**Fig. 2a**, middle). While both dimerization arms were resolved in the EGFR/EREG structure, they displayed significantly different B-factors and were differentially engaged. The dimerization arm engaged with the partially open binding pocket was more dynamic and formed only a single hydrogen bond with the dimerization partner (**Fig. 2b**, **Extended Data Fig. 4**). The disengagement of this dimerization arm represents an intermediate state between the missing dimerization arm of HER3 in our structure, and the fully engaged dimerization arms, both displaying low B-factors, in the symmetric EGFR/EGF ectodomain dimer (**Fig. 2b**).

**Fig. 2.**
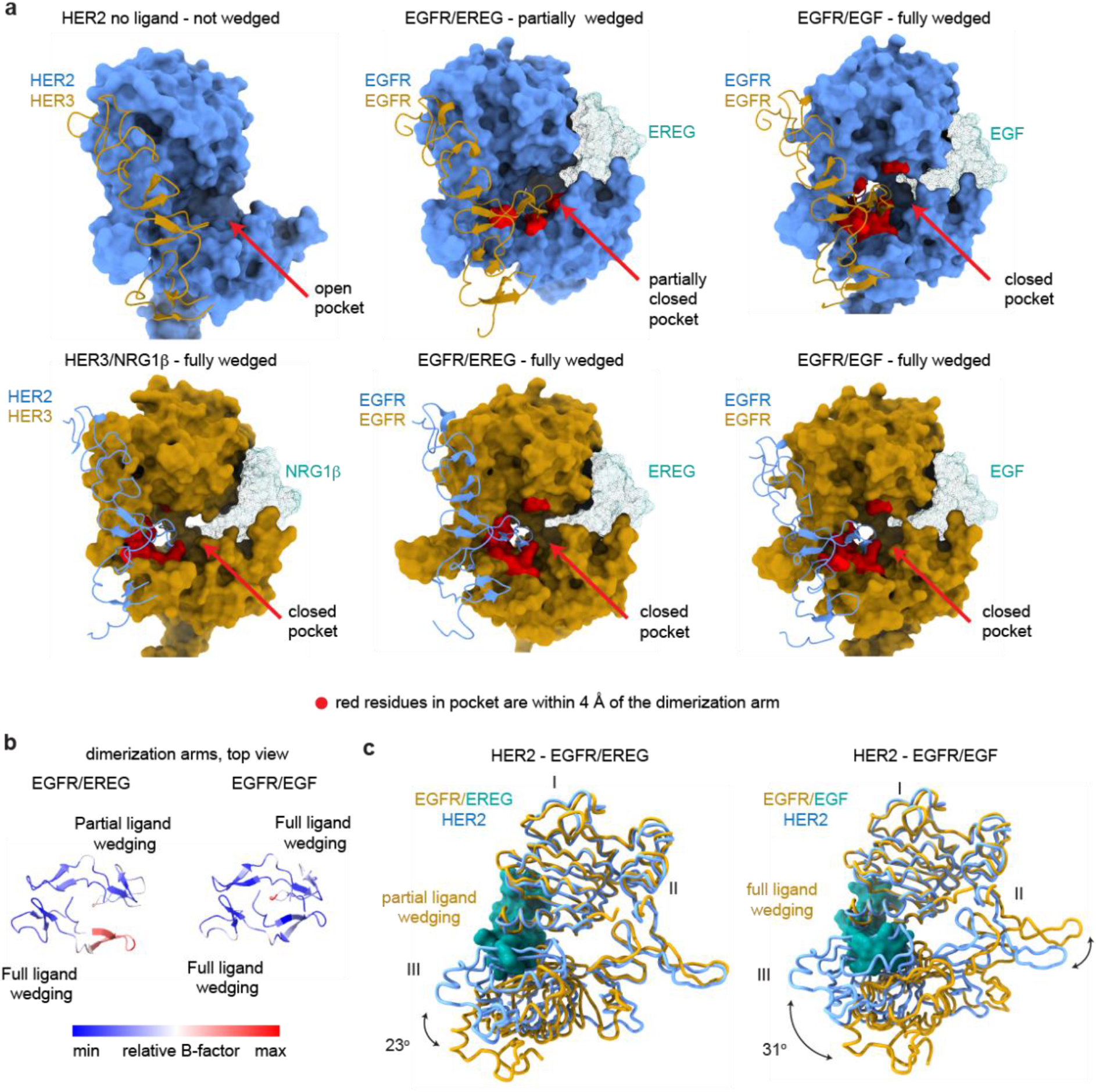
Analysis of liganded HER receptor states reveals an allosteric mechanism of dimerization arm engagement. **a,** Left top panel - an open dimerization arm binding pocket in the ligand-free HER2 does not engage HER3 dimerization arm in the HER2/HER3/NRG1β structure. Left bottom panel – closed binding pocket in HER3 engages the HER2 dimerization arm in the same structure. Middle top panel – a partially closed dimerization arm binding pocket in one of the monomers in the EGFR/EREG structure (PDB: 5WB7) in which the ligand (EREG) is partially-wedged. Middle bottom panel - closed binding pocket in another monomer in the same EGFR/EREG structure in which the ligand is fully-wedged. Right panel top and bottom shows the identical conformations of both EGFR monomers in the EGFR/EGF dimer structure (PDB: 3NJP) in which the ligands (EGF) are fully wedged. Consequently, both dimerization arms are engaged. Residues within 4Å of the dimerization arm are shown in red. **b,** Top view of dimerization arms in the asymmetrically ligand-wedged EGFR/EREG and symmetrically ligand-wedged EGFR/EGF crystal structures indicating different values of B-factors (PDB: 5WB7 and 3NJP, respectively). **c,** Detailed view of domains I-III in the EGFR/EREG or EGFR/EGF crystal structures aligned on HER2 domain I in the structure of the HER2/HER3/NRG1β complex. The EGFR monomer in which the EREG ligand is only partially-wedged is shown. The extent of ligand wedging between domains I and III induces a graded rotation of domain III as compared to domain III of HER2.

What determines the conformation of the dimerization arm-binding pocket? Our analysis points to its previously unappreciated allosteric connection with the ligand binding site. Thus far, two modes of ligand binding to HER receptors have been described: fully-wedged and partially-wedged^32^. These modes are differentiated based on the relative rotation between domains I and III induced by ligand binding. The fully-wedged conformation, as seen in the symmetric EGFR/EGF, EGFR/TGFα and HER4/NRG1β dimer structures, is characterized by a ~31° rotation between domains I and III (the reference point is the angle between domains I and III in HER2). The partially-wedged ligand conformation, as seen in EGFR/EREG and *Drosophila* EGFR/Spitz structures, is associated with a smaller ~23° rotation. These different conformations directly correlate with the state of the dimerization arm-binding pocket on the other side of the receptor (**Fig. 2c**). A fully-wedged ligand results in the formation of a closed high affinity dimerization arm-binding pocket. Consequently, the HER3 monomer, which in our structure has a fully-wedged ligand with the associated large rotation of domains I and III, provides a high affinity pocket for the HER2 dimerization arm. Partial ligand-wedging results only in partial closure of the dimerization arm-binding pocket, and hence the increased dynamics of the dimerization arm presented by the dimerization partner as seen in one monomer of the EREG/EGFR dimer^32^. In HER2, which does not bind a ligand, domains I and III do not undergo a relative rotation, and consequently the dimerization arm-binding pocket is fully open and does not engage the HER3 dimerization arm in our structure (**Fig. 2a, c**).

The conformational coupling between bound ligand and dimerization arm binding pocket explains why solution and structural studies of the extracellular domains of HER2 have been unable to identify homodimers despite HER2 being always in an extended conformation with the dimerization arm exposed^25,26,34^. Our structure shows that because the ligand-free extracellular domain of HER2 does not undergo a necessary rotation between domains I and III, it cannot establish a binding pocket for the partner’s dimerization arm. Therefore, the extracellular domains of HER2 are effectively protected from homo-association. However, HER2 can bind ligand-bound HER receptors, in which the dimerization arm-binding pocket is established and can engage the HER2 dimerization arm.

### The most common oncogenic HER2 variant enhances heterodimerization by stabilizing the dimerization arm of HER3

The most frequent cancer-associated missense mutation in HER2 is localized in the domain II of the extracellular domain and changes serine 310 to a phenylalanine or a tyrosine (S310F/Y). This mutant variant of HER2 found in cancers without HER2 overexpression enhances anchorage-dependent colony formation, HER2-dependent autophosphorylation, and is strongly proliferative^35,36,4^. Interestingly, S310 is located in the direct vicinity of the dimerization arm-binding pocket in HER2. To test if this mutant may influence the interactions between HER2 and HER3, we expressed, purified and reconstituted a nearly full-length HER2-S310F/HER3/NRG1β complex *in vitro*. When compared to the wild type HER2, significantly more HER2-S310F was captured by HER3 immobilized on NRG1β-bound beads (**Extended Data Fig. 5**) suggesting that the mutation substantially stabilizes the HER2/HER3 heterodimer.

We obtained a cryo-EM reconstruction of the extracellular module of the HER2-S310F/HER3/NRG1β complex at 3.1Å resolution (**Fig. 3a**, **Extended Data Fig. 6**). While the structure closely resembles that of the complex containing wild type HER2 with no conformational changes in the HER2 monomer (RMSD: 0.1 Å), remarkably, the HER3 dimerization arm was entirely resolved in the mutant complex (**Fig. 3a,b**). The main stabilizing interaction involves π-π stacking between the introduced phenylalanine at position 310 (HER2-F310) and HER3-Y265 (**Fig. 3c**). This new interaction also positioned the HER3 dimerization arm such that stabilizing polar contacts formed between the HER3-Y265 sidechain hydroxyl group and the backbones of HER2-F290 and HER2-C311 (**Fig. 3c**). The stabilized HER3 dimerization arm increases the total buried surface area at the HER2/HER3 interface (domains I-III) from 1,821 Å^2^ in the wild type complex to 3,054 Å^2^ in the mutant complex, which is even higher than the respective interfaces in structures of the symmetric ligand-bound EGFR and HER4 homodimers (**Extended Data Fig. 4**)^23,24^. We predict that the same mechanism is employed by the HER2 S310Y mutation, which is assumed to form an analogous π-π stacking interaction with HER3 Y265 and similarly stabilize the heterocomplex by engaging the HER3 dimerization arm. Thus, the most common HER2 oncogenic mutations act by stabilizing interactions with the HER3 dimerization arm and compensate for the inability of HER2 to undergo a needed rotation between domains I and III.

**Fig. 3.**
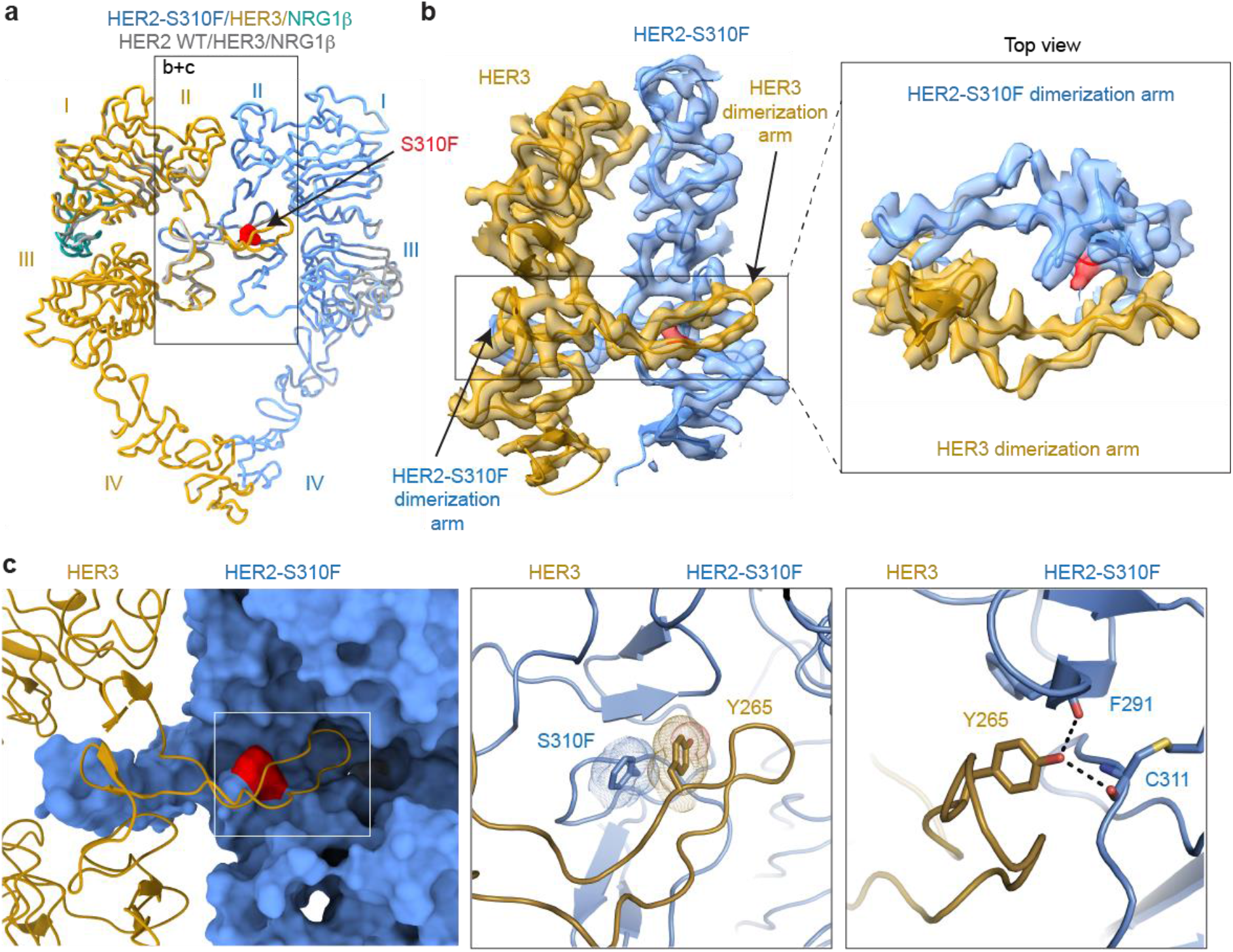
HER2 oncogenic mutation S310F stabilizes the dimerization arm of HER3. **a,** Cryo-EM structure of HER2-S310F/HER3/NRG1β complex overlayed on HER2/HER3/NRG1β. The HER2-S310F mutation is shown in red. **b,** Cryo-EM map and model zoomed in on domains II depict a resolved HER3 dimerization arm in the HER2-S310F/HER3/NRG1β complex. Inset shows a top-down view of the HER2 and HER3 dimerization arms. **c,** Left panel, HER2-S310F monomer, shown in surface representation, pins the HER3 dimerization arm, shown as cartoon, in the HER2 dimerization arm-binding pocket despite its inability to close in the ligand-less HER2. Middle panel, HER2-S310F forms a π-π interaction with HER3 Y265 that stabilizes the dimerization arm. Right panel, polar contacts (dotted lines) between HER3 Y265 and the backbone residues of HER2 - F291 and C311.

### The HER2/HER3 heterodimer accommodates trastuzumab and pertuzumab Fabs

The clinically-approved HER2-targeting monoclonal antibodies, trastuzumab and pertuzumab, target domains IV and II, respectively^25,37^. To assess if these therapeutic agents also bind the HER2/HER3 heterocomplex, we incubated NRG1β-stabilized HER2/HER3 heterodimers with an excess of trastuzumab or pertuzumab Fab, followed by HER2 affinity purification and evaluation of bound HER3 and Fab levels. We found that neither trastuzumab nor pertuzumab interfered with HER2/HER3 heterodimerization and could be found associated with the receptor dimers as ternary complexes (**Fig. 4a**).

**Fig. 4.**
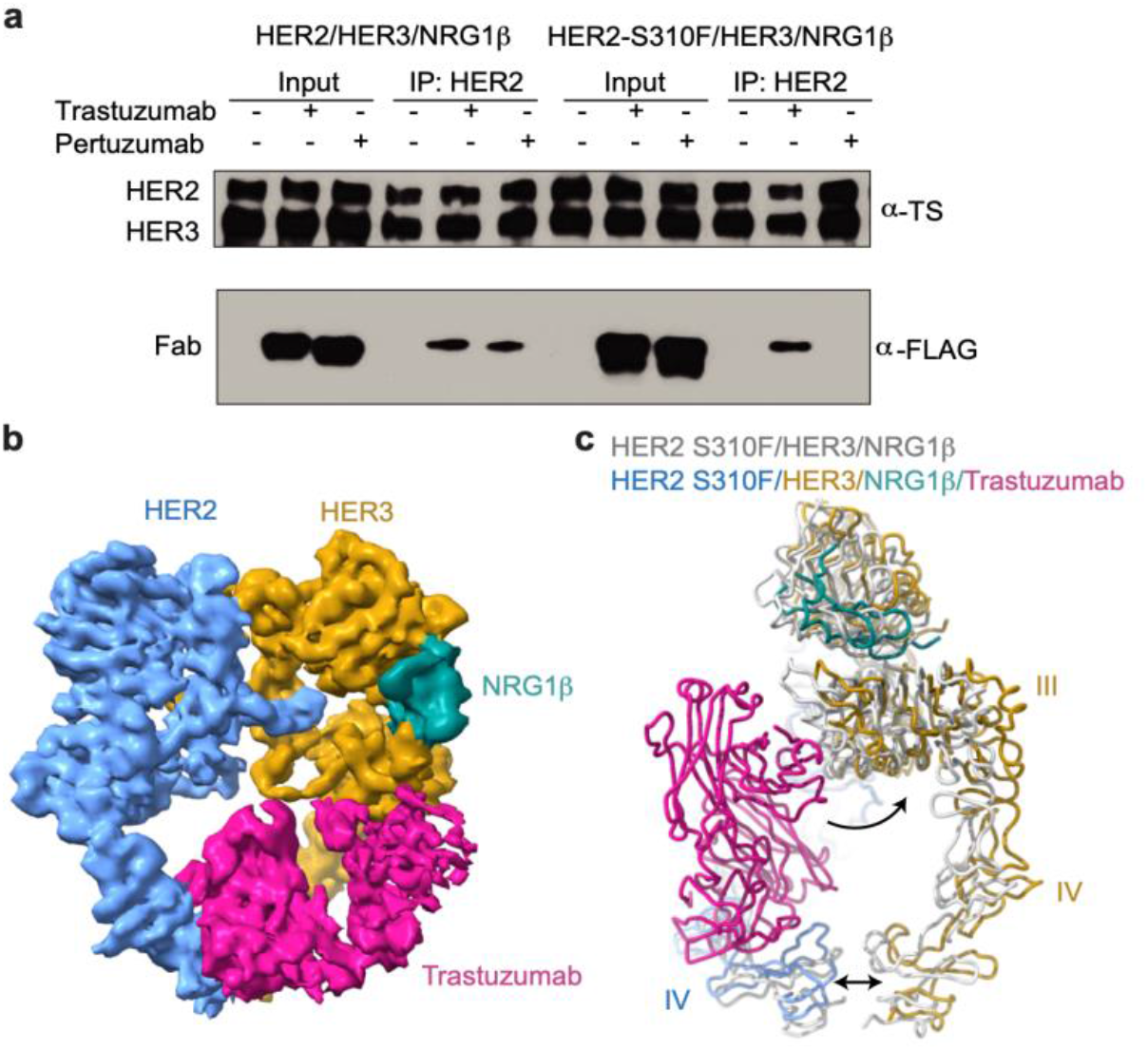
The HER2/HER3/NRG1β structure accommodates trastuzumab binding. **a,** Representative Western blot of heterodimer pulldowns in the presence of a two-fold molar excess of pertuzumab and trastuzumab FLAG-tagged Fabs. Pre-formed HER2/HER3/NRG1β heterocomplex binds both Fabs whereas HER2-S310F/HER3/NRG1β only binds the trastuzumab Fab. TS – Twin Strep tag (present on both HER2 and HER3). **b,**5Å lowpass filtered density of the HER2-S310F/HER3/NRG1β heterocomplex bound to trastuzumab Fab. **c,** Ribbon overlay of the HER2-S310F/HER3/NRG1β heterocomplex with (multi-color) and without (light grey) trastuzumab Fab. The Fab pushes HER3 back relative to HER2 (curved arrow) and spreads domains IV further apart (double-headed arrow).

The ability of the HER2/HER3 complex to associate with trastuzumab is rationalized by our cryo-EM reconstruction of the trastuzumab Fab bound to the HER2-S310F/HER3 heterodimer that we obtained at 3.5Å resolution (**Fig. 4b**, **Extended Data Fig. 7**). Our structure shows that the HER2/HER3 heterodimer rearranges in multiple regions to accommodate trastuzumab binding to domains II and IV. Specifically, domain IV of HER2 moves away from HER3 as a rigid body with the variable domains of trastuzumab. In addition, HER3 rotates in relation to HER2 to resolve a steric clash between HER3 domain III and the constant domains of the trastuzumab Fab (**Extended Data Fig. 8**, **Fig. 4c**). Thus, minor structural rearrangements and the previously noted flexibility in domain IV underlie trastuzumab binding to the HER2/HER3 heterodimer.

Our observation that pertuzumab binds to the HER2/HER3/NRG1β complex is less straightforward albeit possible to rationalize. Pertuzumab docked into the structure of the HER2/HER3/NRG1β heterodimer clashes with the domain II of HER3, directly blocking the extracellular domain dimerization interface (**Extended Data Fig. 8**). With pertuzumab bound, the extracellular domains of HER2 and HER3 are unlikely to interact. Thus, the fact that pertuzumab still does not interfere with HER2/HER3 dimers emphasizes the important role the intracellular receptor domains play in stabilizing the interaction between HER2 and HER3. We were unable to obtain high resolution reconstruction of the pertuzumab-bound extracellular domain module of the HER2/HER3/NRG1β heterocomplex which supports the notion that pertuzumab binding increases its dynamics.

Similar to the wild type HER2/HER3/NRG1β complex, the presence of trastuzumab or pertuzumab did not affect assembly of the mutant HER2-S310F/HER3/NRG1β complex (**Fig. 4a**). However, in a stark contrast to the wild type complex, the mutant complex did not bind to pertuzumab. This could be explained by direct interference of the S310 mutation with the Fab binding (**Extended Data Fig. 8).** It is also possible that the HER2 epitope recognized by pertuzumab is occluded in the mutant complex due to the enhanced extracellular domain dimerization interface (**Extended Data Fig.8**). Thus, our work suggests that pertuzumab may be less effective than trastuzumab at targeting cancers driven by HER2-S310F/Y.

## Discussion

The HER2/HER3/NRG1β captures two obligate heterodimeric HER receptors in an active state revealing the previously unseen ligand-bound extended state of HER3 engaging the ligand-free HER2. Our structure expands on the repertoire of ligand-bound states of human HER receptors by representing the first heterodimeric complex of HER extracellular domains and the first singly-liganded human HER dimer. A singly-liganded dimer was previously seen only in *Drosophila* EGFR^38^, and was suggested to represent a more stable active complex than a doubly liganded one. To the contrary, we observe that the singly liganded HER2/HER3 complex is more dynamic compared to other solved HER receptor extracellular domain dimer structures, to the extent that one dimerization arm is not even engaged at the dimerization interface. The increased dynamics are rationalized by allosteric coupling between the growth factor-binding pocket and the dimerization arm-binding pocket within the same receptor molecule. The resulting closure of the ligand binding pocket during the transition from a fully-wedged ligand bound state, as observed in HER3, to the apo state, as seen in HER2, leads to a gradual decrease in the ability of a receptor to interact with the dimerization arm of its partner. As shown in our structure, ligand-less HER2 cannot in fact bind the dimerization arm of HER3. Such a destabilized dimerization arm interface has been previously observed in molecular dynamics simulations of the putative EGFR-HER2 ECD dimer, underlining the generalizability of our findings to other HER2-containing heterocomplexes^39^. Furthermore, this allosteric model posits that ligands which bind HER receptors in partially-wedged conformations will increase dynamics at the dimerization interface. Indeed, this is the case for the low affinity ligands of EGFR, like EREG and epigen (EPGN)^32^.

Our findings demonstrate that HER2 does not undergo a significant conformational change when it complexes with HER3, suggesting that the constitutively extended state, previously deemed as autoinhibited^27^, is in fact dimerization-competent. However, in order to dimerize, HER2 relies on co-receptors to engage its dimerization arm while HER2 itself cannot reciprocate. This is likely because the constitutively closed growth factor binding pocket in HER2 leads to the loss of an allosteric connection to its own dimerization arm binding pocket. Thus, the HER2 extracellular domain is specifically autoinhibited towards self-association but receptive to heterodimerization.

Our structure of the mutant HER2-S310F/HER3 heterocomplex reveals how cancer subverts the intrinsic dynamics at the HER2-containing extracellular heterodimer interface leading to an aberrantly stabilized heterodimer, a fundamentally new mechanism of driving aberrant HER2 signaling. We envision that such mutations would cooperate with growth factor binding to HER2 dimerization partners (like HER3) to promote the dimerization-arm exposed extended conformation of the partner receptor which would in turn be further stabilized by the mutant HER2. The notable dynamics at the interface likely also accounts for the ability of the wild-type HER2/HER3/NRG1β heterocomplex to accommodate both trastuzumab and pertuzumab binding *in vitro*. Our structure of the trastuzumab-bound HER2-S310F/HER3/NRG1β reveals how the heterodimer overcomes a steric clash to accommodate the trastuzumab Fab. Finally, our results show that the HER2-S310F/HER3 complex resists pertuzumab binding, which is of therapeutic significance. The HER2/HER3 structures presented here provide a long-awaited platform for the rational design of therapeutics and biomarkers specific to the active states of these receptors and their complexes.

## AUTHOR CONTRIBUTIONS

N.J. conceived of project and D.D., R.T., K.A.V., and N.J. designed the research approach. D.D. and R.T. performed all expression and purification, electron microscopy imaging and processing, structural modelling, and *in vitro* experiments. T.M.T. provided initial receptor expression constructs. F.W. and D.A. provided holey carbon graphene-oxide grids for cryo-EM. D.D., R.T, K.A.V., and N.J. wrote the manuscript.

## COMPETING INTERESTS

N.J. is a member of the SAB and a shareholder of Turning Point Therapeutics, SUDO Biosciences and Type6 Therapeutics. The Jura laboratory has received sponsored research support from Genentech. Other authors do not declare competing interests.

## ACKNOWLEDGMENTS

We thank A. Manglik for advice on receptor expression, and D. Bulkley, G. Gilbert, E. Tse, and Z. Yu from the UCSF EM facility. We also thank M. Moasser, S. Seshagiri, and members of the Verba and Jura labs for their helpful discussions, and E. Linossi and H. Torosyan for critical comments on the manuscript. We would also like to acknowledge D. Suveges for his experimental contributions to the generation of our first HER expression constructs. This work was funded through UCSF Program for Breakthrough Biomedical Research to K.V. and N.J., NIH/NIGMS R35-GM139636 to N.J., Genentech research grant to N.J., DFG German Research Foundation GZ: TR 1668/1-1 to R.T. and NIH/NCI 1F30CA247147 to D.D.

## MATERIALS AND METHODS

### NRG1β expression and purification

An HRV-3C cleavable Thyrodoxin A (TrxA) was fused to the EGF-like domain of NRG1β (residues 177-236, NRG1 isoform 6 (UniProt: Q02297-6; numbering includes propeptide) with C-terminal Flag and 6x-Histidine tags and subsequently cloned into a p32A vector. The TrxA-3C-NRG1β-Flag-6xHis construct was transformed into Origami *E. coli*, grown at 37 °C in Terrific Broth in large scale culture until an OD of ~1.0 - 1.5, and induced with 1 mM Isopropyl β-d-1-thiogalactopyranoside (IPTG) overnight at room temperature. Cells were harvested the next day, pelleted, flash frozen, and stored until purification. For the purification, cells were resuspended in NRG lysis buffer (50 mM Tris-HCl pH 7.4, 150 mM NaCl, 1 mM phenylmethylsulfonyl fluoride (PMSF), and protease inhibitors (eOComplete, Roche)) and sonicated until thoroughly lysed. Lysate was then clarified through ultracentrifugation, syringe filtered through 0.44 μm filters and incubated with Ni-NTA resin overnight. The beads were washed by gravity through 20 column volumes (CVs) of NRG wash buffer (50 mM Tris-HCl pH 7.4, 150 mM NaCl) containing 20 mM imidazole, then 10 CVs of NRG wash buffer containing 50 mM imidazole, and finally eluted with 3 CVs of NRG wash buffer containing 300 mM imidazole. Imidazole in the eluate was reduced < 30 mM over a 10K concentrator and subsequent dilution with NRG wash buffer. The eluate was cleaved overnight with 3C protease at 4 °C. To remove cleaved TrxA, the elution was again applied on equilibrated Ni-NTA resin, incubated, washed, and eluted as described previously. The elution containing NRG1β was concentrated with a 3K cutoff and applied on an S200 10/300 increase column (GE Healthcare). Protein from the major peak was stored in aliquots at −80 °C for subsequent receptor purifications. Yields ranged from 5-10 mg/L of culture.

### Trastuzumab and pertuzumab Fab expression and purification

The Fragment antigen binding (Fab) heavy chain and light chain sequences encoding trastuzumab and pertuzumab were inserted into the pSVF4 vector. For each Fab, a 1x Flag tag was inserted after the heavy chain constant domain and a 6x-Histidine tag was inserted after the light chain constant domain. Constructs were transformed into BL21 gold *E. coli* and scaled up to 6L in 2xYT media under Ampicillin antibiotic selection. Cultures were grown at 37 °C until OD of 0.8 - 1.0, induced with 1 mM IPTG for 6 hrs at 37 °C, harvested by centrifugation, and stored at −80 °C. Cells were resuspended in 100 ml of lysis buffer (20 mM sodium phosphate pH 7.4, 500 mM NaCl with DNase I (Roche), 0.5 mM MgCl_2_ and 1 mM PMSF). Cells were sonicated until fully lysed and resulting lysate was incubated at 65 °C for 30 minutes. The lysate was cooled on ice and spun down at 40,000 rpm for 60 minutes at 4 °C. The clarified lysate was loaded onto a Protein A column equilibrated in Buffer A (20 mM sodium phosphate pH 7.4, 500 mM NaCl), washed with 10 column volumes of Buffer A, and eluted in 100 mM acetic acid by fractionation into neutralizing buffer containing 20 mM Tris-HCl pH 9.0, 150 mM NaCl. Immediately following Protein A purification, eluent was concentrated and loaded onto a Superdex 200 10/300 Increase column (GE Healthcare) equilibrated in a buffer containing 50 mM Tris-HCl pH 7.4, 150 mM NaCl. Fractions corresponding to Fab were pooled and stored at 4 °C until needed.

### Near full-length receptor expression

Human HER2 with a C-terminal tail truncation (Δ1030–1255) followed by maltose binding protein (MBP) and twin-strep tags was cloned into pFastBac with a CMV promoter. A single point mutation in the HER2 kinase domain, G778D, which confers Hsp90 independence^40^, was introduced to improve yields. Human HER3 with a C-terminal tail truncation (Δ1023 – 1342) followed by a twin-strep tag was cloned in pFastBac with a CMV promoter. Two oncogenic mutations that stabilize the asymmetric kinase domain dimer, Q709R and E928G, were introduced to further improve heterodimer yields^11,17^. The HER2 and HER3 constructs were each transfected into 60 ml of Expi293 mammalian suspension (Life Technologies) cells cultured to 4×10^6^ cells/ml at 37 °C, 8% CO_2_ following the standard expression protocol. 10 mM canertinib in DMSO was added 16 - 18 hrs post-transfection to a final concentration of 10 μM along with ExpiFectamine 293 Transfection Kit enahncers 1 and 2. Cells were harvested, flash frozen, and stored at −80 °C 24 hrs after the addition of enhancers. The same procedure was followed for HER2 in the presence and absence of S310F mutation.

### Heterodimer purification

Cells were resuspended with the lysis buffer (50 mM Tris-HCl pH 7.4, 150 mM NaCl, 1 mM NaVO_3_, 1 mM NaF, 1 mM EDTA, protease inhibitors (eOComplete, Roche), DNAse I, and 1% DDM (Inalco)) and lysed for 2 hrs by gentle rocking at 4 °C. Lysate was clarified by centrifugation at 4,000g for 20 min at 4 °C. Purified EGF-like domain of NRG1β was incubated with G1 Flag Resin (Genscript) for 1 hr at 4 °C and serially washed 3x with Buffer A (50 mM Tris-HCl pH 7.4, 150 mM NaCl). Clarified HER2 and HER3 receptor lysates were mixed and incubated O/N in batch mode at 4 °C with NRG1β Flag beads. NRG1β Flag beads were serially 3x washed with Buffer A containing 0.5 mM DDM (Anatrace) and eluted with Buffer A containing 0.5 mM DDM and 250 μg/ml of Flag peptide (SinoBiological). The eluate was then applied to amylose resin in batch mode for 2 hrs, washed serially 3x with Buffer B (50 mM HEPES pH 7.4, 150 mM NaCl) containing 0.5 mM DDM and eluted with amylose elution buffer (Buffer B containing 0.5 mM DDM and 10 mM maltose) O/N at 4 °C. The eluate was concentrated to 0.4 ml with a 100-kDa concentrator (Amicon) and mildly crosslinked in 0.2% glutaraldehyde for 40 min on ice. The sample was loaded on a Superose6 10/300 (GE Healthcare) pre-equilibrated with Buffer A containing 0.5 mM DDM and 0.5 ml fractions were collected. Peak fractions corresponding to heterodimer sample were pooled, concentrated down to ~20 μl with a 100-kDa concentrator, and flash frozen for grid preparation. The same purification protocol was followed for HER2-S310F/HER3 heterocomplex. The HER2-S310F/HER3 + trastuzumab Fab complex sample was generated by incubating a 5x molar excess of Fab with the heterocomplex prior to crosslinking, gel filtration, and imaging.

### Electron microscopy sample preparation and imaging

For negative stain EM, fractions corresponding to heterodimer were applied to negatively glow-discharged carbon coated copper grids, stained with 0.75% uranyl-formate, and imaged on an FEI-Tecnai T12 with an 4k CCD camera (Gatan). The resulting negative stain micrographs were assessed for particle homogeneity and particle density. This analysis was used to determine the target concentration for cryo-EM with graphene oxide grids which typically require 2-5x negative stain concentrations.

For cryo-EM, 3 μl of purified and concentrated heterodimer sample (as empirically determined by negative stain) was applied to graphene-oxide coated Quantifoil R1.2/1.3 300 mesh Au holey-carbon grids prepared as previously described^30^, blotted using a Vitrobot Mark IV (FEI) and plunge frozen in liquid ethane (no glow discharge, 30 second wait time, room temperature, 100% humidity, 4-8 seconds blot time, 0 blot force).

Grids were imaged on a 300-keV Titan Krios (FEI) with a K3 direct electron detector (Gatan) and a BioQuantum energy filter (Gatan). Data for HER2/HER3/NRG1β and HER2-S310F/HER3/NRG1β were collected in super-resolution mode at a physical pixel size of 0.835Å/pix with a dose rate of 8.0 e^-^ per pixel per second (operated in CDS mode). Images were recorded with a 5.9 s exposure over 118 frames with a dose rate of 0.57 e^-^/Å^2^/frame. Data for HER2-S310F/HER3/NRG1β with trastuzumab Fab were collected in super-resolution pixel mode at a physical pixel size of 0.834Å/pix with a dose rate of 8.0 e^-^ per pixel per second. Images were recorded in 6 s exposures over 120 subframes with a dose rate of 0.55 e^-^/Å^2^/frame.

### Image processing and 3D reconstruction

Raw movies were corrected for motion and radiation damage with MotionCor2^41^ and the resulting sums were imported in CryoSPARC2^42^. To account for the reduced GO coverage with detergent sample, all datasets underwent strict micrograph curation with a final yield of ~40-50% of the collected micrograph stack. Micrograph CTF parameters were estimated with the patch CTF job in CryoSPARC2. Particles were picked through template picking with the extracellular domain volume of HER4 (PDB ID: 3U7U) low-pass filtered to 25Å, the resulting picks were extracted with 2x Fourier cropping and subjected to iterative rounds of *ab initio* and heterogeneous refinements. Once reasonable reconstructions were obtained (as judged by the FSC (Fourier Shell Correlation) curves shape), unbinned particles were re-extracted and run through subsequent rounds of heterogeneous and non-uniform refinements to achieve reconstructions with the highest resolution. The final reconstruction of HER2/HER3/NRG1β used for model building included 123,173 particles with C1 symmetry and resulted in an overall resolution of 2.9Å by Gold Standard-Fourier Shell Correlation (GS-FSC) cutoff of 0.143. The final reconstruction of HER2-S310F/HER3/NRG1β used for model building included 99,755 particles with C1 symmetry and attained a GS-FSC resolution of 3.1Å.

HER2-S310F/HER3/NRG1β/Trastruzumab Fab data set was initially processed as above in CryoSPARC2. To address incomplete Fab occupancy, a stack containing 330,000 particles was imported into RELION3^43^ and subclassified through skip-align classification. Particles classified into reconstructions without Fab density were removed from the particle stack. A final particle stack from RELION3 containing 243,376 particles was re-imported into CryoSPARC2 and subjected to non-uniform refinement to produce a reconstruction with a final resolution of 3.45Å.

Each map was assessed for local and directional resolutions through ResMap^44^ and 3DFSC^45^ server respectively. For all reconstructions, extracellular domains I-III achieved the highest local resolutions (~3Å) while that of domain IV varied from 4-8Å suggesting that a high degree of flexibility exists closer to the transmembrane domains. All reconstructions achieved a sphericity > 0.9.

To recover micelle and sub-micelle densities, 2x binned particle stacks for HER2/HER3/NRG1β and HER2-S310F/HER3/NRG1β were imported into RELION3 and further 3D-classified. Particles classified into 3D classes with substantial micelle densities were re-extracted with shifted coordinates (PyEM^46^) on the center of the micelle and refined. Resulting reconstructions featured convincing sub-micelle density with volumes large enough to accommodate transmembrane domains and kinases.

### Model refinement and validation

An initial model was generated by docking HER2 (PDB ID: 1N8Z) with a homology model of liganded HER3 from its closest homolog, HER4, in SwissProt (PDB ID: 3U7U) into the HER2-S310F/HER3/NRG1β map. Given the substantial variation in domain IV local resolution, domain IV was truncated from the model and domains I-III were iteratively rebuilt in Rosetta^47^. Top scoring models were selected and further edited in Coot^48^ and ISOLDE^49^. Domains IV were then placed into the model (HER2 PDB ID: 6OGE, HER3 PDB ID: 1M6B) and fit into the density with a FastRelax Rosetta protocol in torsion space. For HER2-S310F/HER3/NRG1β + trastuzumab Fab, the Fab (PDB ID: 6OGE) was torsion relaxed with the HER2-S310F/HER3/NRG1β model in Rosetta.

For glycan building, glycans were initially manually placed into the density in Chimera^50^ and then were refined with the Rosetta glycan refinement protocol^51^. Model statistics were routinely assessed in PHENIX^52^ and glycan geometries were cross validated in Privateer^53^. All structures were deposited into the EMDB and Protein Data Bank (PDB).

### Small-scale heterodimer pulldowns and Western Blot

Tagged HER2 and HER3 expression constructs were co-transfected into 2 ml cultures of Expi293 cells as described above. Cell pellets were lysed in 1 ml lysis buffer and clarified lysates were subjected to NRG-pulldown and eluted in 250 μl Flag elution buffer as described above. The extent of heterodimer formation was assessed by Western blot. Samples were boiled in SDS-loading buffer at 95 °C for 5 min, run on 4-15%acrylamide gels and transferred onto PVDF membranes. Membranes were blocked in 3% BSA in TBS with 0.1% Tween (TBST) overnight and incubated with Step-Tactin HRP (IBA, 1:5000) in TBST + 3% BSA for 1 hr at room temperature. Membranes were washed 5x with TBST and signal was detected using ECL Western Blotting detection reagent (GE) or ECL prime (VWR).

### Trastuzumab and pertuzumab pulldown assay

Wild-type HER2 and S310F HER2 heterodimeric complexes with HER3 were expressed and purified as described above until elution from amylose resin with the exception that amylose wash and elution buffers contained 50 mM Tris-HCl pH 7.4 instead of 50 mM HEPES pH 7.4. Eluates were concentrated to 100 μl and maltose was removed via buffer exchange using 7 MWCO Zeba spin desalting columns. 70 nM heterodimer solutions were each incubated with 1 and 10x molar ratios for 30 min and bound to amylose resin overnight. Complexes were eluted as described above and complex formation with trastuzumab and pertuzumab were assessed by Western Blot.

## DATA AVAILABILITY STATEMENT

The data that support the findings of this study are available from the corresponding author upon request. 3D cryo-EM density maps have been deposited in the Electron Microscopy Data Bank under the accession number EMD-23916, EMD-23917, and EMD-23918. Atomic coordinates for the atomic models have been deposited in the Protein Data Bank under the accession number 7MN5, 7MN6, and 7MN8.

## REFERENCES

1 Sliwkowski, M. X. et al. Coexpression of erbB2 and erbB3 proteins reconstitutes a high affinity receptor for heregulin. J Biol Chem 269, 14661–14665 (1994).

2 Wallasch, C. et al. Heregulin-dependent regulation of HER2/neu oncogenic signaling by heterodimerization with HER3. The EMBO Journal 14, 4267–4275 (1995).

3 Moasser, M. M. The oncogene HER2: its signaling and transforming functions and its role in human cancer pathogenesis. Oncogene 26, 6469–6487, doi:10.1038/sj.onc.1210477 (2007).

4 Network, T. C. G. A. Comprehensive Molecular Portraits fo Human Breast Tumors. Nature 490, 61–70, doi:10.1038/nature11412 (2012).

5 Lee, J. W. et al. Somatic mutations of ERBB2 kinase domain in gastric, colorectal, and breast carcinomas. Clin Cancer Res 12, 57–61, doi:10.1158/1078-0432.CCR-05-0976 (2006).

6 Bose, R. et al. Activating HER2 mutations in HER2 gene amplification negative breast cancer. Cancer Discov 3, 224–237, doi:10.1158/2159-8290.CD-12-0349 (2013).

7 C., L. et al. c-erbB-2 Oncoprotein Expression in Primary and Advanced Breast Cancer. British Journal of Cancer 63, 439–443 (1991).

8 Slamon, D. J. et al. Human breast cancer: correlation of relapse and survival with amplification of the HER-2/neu oncogene. Science 235, 177–182 (1987).

9 Peters, S. & Zimmermann, S. Targeted therapy in NSCLC driven by HER2 insertions. Transl Lung Cancer Res 3, 84–88, doi:10.3978/j.issn.2218-6751.2014.02.06 (2014).

10 Connell, C. M. & Doherty, G. J. Activating HER2 mutations as emerging targets in multiple solid cancers. ESMO Open 2, e000279, doi:10.1136/esmoopen-2017-000279 (2017).

11 Jaiswal, B. S. et al. Oncogenic ERBB3 mutations in human cancers. Cancer Cell 23, 603–617, doi:10.1016/j.ccr.2013.04.012 (2013).

12 Sergina, N. V. et al. Escape from HER-family tyrosine kinase inhibitor therapy by the kinase-inactive HER3. Nature 445, 437–441, doi:10.1038/nature05474 (2007).

13 Garrett, J. T. et al. Transcriptional and posttranslational up-regulation of HER3 (ErbB3) compensates for inhibition of the HER2 tyrosine kinase. Proc Natl Acad Sci U S A 108, 5021–5026, doi:10.1073/pnas.1016140108 (2011).

14 Sierke, S. L., Cheng, K., Kim, H. & Koland, J. G. Biochemical characterization of the protein tyrosine kianse domain of the ErbB3 (HER3) receptor protein. Biochem J 322, 757–763 (1997).

15 Jura, N., Shan, Y., Cao, X., Shaw, D. E. & Kuriyan, J. Structural analysis of the catalytically inactive kinase domain of the human EGF receptor 3. Proc Natl Acad Sci U S A 106, 21608–21613, doi:10.1073/pnas.0912101106 (2009).

16 Zhang, X., Gureasko, J., Shen, K., Cole, P. A. & Kuriyan, J. An allosteric mechanism for activation of the kinase domain of epidermal growth factor receptor. Cell 125, 1137–1149, doi:10.1016/j.cell.2006.05.013 (2006).

17 Littlefield, P. et al. Structural analysis of the EGFR/HER3 heterodimer reveals the molecular basis for activating HER3 mutations. Science Signaling 7 (2015).

18 Ferguson, K. et al. EGF Activates Its Receptor by Removing Interactions with Autoinhibit Ectodomain Dimerization. Molecular Cell 11, 507–517 (2003).

19 Bouyain, S., Longo, P. A., Li, S., Ferguson, K. M. & Leahy, D. J. The extracellular region of ErbB4 adopts a tethered conformation in the absence of ligand. Proc Natl Acad Sci U S A 102, 15024–15029, doi:10.1073/pnas.0507591102 (2005).

20 Cho, H.-S. & Leahy, D. J. Structure of the Extracellular Region of HER3 Reveals an Interdomain Tether. Science 297, 1330–1333 (2002).

21 Garrett, J. T. et al. Crystal Structure of a Truncated Epidermal Growth Factor Receptor Extracellular Domain Bound to Transforming Growth Factor alpha. Cell 110, 763–773 (2002).

22 Ogiso, H. et al. Crystal Structure of the Complex of Human Epidermal Growth Factor and Receptor Extracellular Domains. Cell 110, 775–787 (2002).

23 Liu, P. et al. A single ligand is sufficient to activate EGFR dimers. Proc Natl Acad Sci U S A 109, 10861–10866, doi:10.1073/pnas.1201114109 (2012).

24 Lu, C. et al. Structural evidence for loose linkage between ligand binding and kinase activation in the epidermal growth factor receptor. Mol Cell Biol 30, 5432–5443, doi:10.1128/MCB.00742-10 (2010).

25 Cho, H.-S. et al. Structure of the extracellular region of HER2 alone and in complex with the Herceptin Fab. Nature 421, 756–760, doi:10.1038/nature01423 (2003).

26 Hao, Y., Yu, X., Bai, Y., McBride, H. J. & Huang, X. Cryo-EM Structure of HER2-trastuzumab-pertuzumab complex. PLoS One 14, e0216095, doi:10.1371/journal.pone.0216095 (2019).

27 Alvarado, D., Klein, D. E. & Lemmon, M. A. ErbB2 resembles an autoinhibited invertebrate epidermal growth factor receptor. Nature 461, 287–291, doi:10.1038/nature08297 (2009).

28 Ferguson, K. M., Darling, P. J., Mohan, M. J., Macatee, T. L. & Lemmon, M. A. Extracellular domains drive homo-but not hetero-dimerization of erbB receptors. The EMBO Journal 19, 4632–4643 (2000).

29 Fry, D. et al. Specific, irreversible inactivation of the epideral growth factor receptor and erbB2, by a new class of tyrosine kianse inhibitor. Proc Natl Acad Sci U S A 95, 12022–12027 (1998).

30 Palovcak, E. et al. A simple and robust procedure for preparing graphene-oxide cryo-EM grids. J Struct Biol 204, 80–84, doi:10.1016/j.jsb.2018.07.007 (2018).

31 Wang, F. et al. General and robust covalently linked graphene oxide affinity grids for high-resolution cryo-EM. Proc Natl Acad Sci U S A 117, 24269–24273, doi:10.1073/pnas.2009707117 (2020).

32 Freed, D. M. et al. EGFR Ligands Differentially Stabilize Receptor Dimers to Specify Signaling Kinetics. Cell 171, 683–695 e618, doi:10.1016/j.cell.2017.09.017 (2017).

33 Huang, Y. et al. A structural mechanism for the generation of biased agonism in the epidermal growth factor receptor. bioRxiv, doi:10.1101/2020.12.08.417006 (2020).

34 Garrett, J. T. et al. The Crystal Structure of a Truncated ErbB2 Ectodomain Reveals an Active Conformation, Poised to Interact with Other ErbB Receptors. Molecular Cell 11, 495–505 (2003).

35 Greulich, H. et al. Functional analysis of receptor tyrosine kinase mutations in lung cancer identifies oncogenic extracellular domain mutations of ERBB2. Proc Natl Acad Sci U S A 109, 14476–14481, doi:10.1073/pnas.1203201109 (2012).

36 Wang, T. et al. HER2 somatic mutations are associated with poor survival in HER2-negative breast cancers. Cancer Sci 108, 671–677, doi:10.1111/cas.13182 (2017).

37 Franklin, M. C. et al. Insights into ErbB signaling from the structure of the ErbB2-pertuzumab complex. Cancer Cell 5, 317–328 (2004).

38 Alvarado, D., Klein, D. E. & Lemmon, M. A. Structural basis for negative cooperativity in growth factor binding to an EGF receptor. Cell 142, 568–579, doi:10.1016/j.cell.2010.07.015 (2010).

39 Arkhipov, A., Shan, Y., Kim, E. T., Dror, R. O. & Shaw, D. E. Her2 activation mechanism reflects evolutionary preservation of asymmetric ectodomain dimers in the human EGFR family. Elife 2, e00708, doi:10.7554/eLife.00708 (2013).

40 Xu, W. et al. Surface charge and hydrophobicity determine ErbB2 binding to the Hsp90 chaperone complex. Nat Struct Mol Biol 12, 120–126, doi:10.1038/nsmb885 (2005).

41 Zheng, S. Q. et al. MotionCor2: anisotropic correction of beam-induced motion for improved cryo-electron microscopy. Nat Methods 14, 331–332, doi:10.1038/nmeth.4193 (2017).

42 Punjani, A., Rubinstein, J. L., Fleet, D. J. & Brubaker, M. A. cryoSPARC: algorithms for rapid unsupervised cryo-EM structure determination. Nat Methods 14, 290–296, doi:10.1038/nmeth.4169 (2017).

43 Scheres, S. H. RELION: implementation of a Bayesian approach to cryo-EM structure determination. J Struct Biol 180, 519–530, doi:10.1016/j.jsb.2012.09.006 (2012).

44 Kucukelbir, A., Sigworth, F. & Tagare, H. Quantifying the local resolution of cryo-EM density maps. Nature Methods 11 (2014).

45 Tan, Y. Z. et al. Addressing preferred specimen orientation in single-particle cryo-EM through tilting. Nat Methods 14, 793–796, doi:10.1038/nmeth.4347 (2017).

46 Asarnow, D., Palovcak, E. & Cheng, Y. UCSF pyem v0.5.. Zenodo (2019).

47 DiMaio, F. et al. Atomic-accuracy models from 4.5-A cryo-electron microscopy data with density-guided iterative local refinement. Nat Methods 12, 361–365, doi:10.1038/nmeth.3286 (2015).

48 Emsley, P., Lohkamp, B., Scott, W. G. & Cowtan, K. Features and development of Coot. Acta Crystallogr D Biol Crystallogr 66, 486–501, doi:10.1107/S0907444910007493 (2010).

49 Croll, T. I. ISOLDE: a physically realistic environment for model building into low-resolution electron-density maps. Acta Crystallogr D Struct Biol 74, 519–530, doi:10.1107/S2059798318002425 (2018).

50 Pettersen, E. F. et al. UCSF Chimera--a visualization system for exploratory research and analysis. J Comput Chem 25, 1605–1612, doi:10.1002/jcc.20084 (2004).

51 Frenz, B. et al. Automatically Fixing Errors in Glycoprotein Structures with Rosetta. Structure 27, 134–139 e133, doi:10.1016/j.str.2018.09.006 (2019).

52 Adams, P. D. et al. PHENIX: a comprehensive Python-based system for macromolecular structure solution. Acta Crystallogr D Biol Crystallogr 66, 213–221, doi:10.1107/S0907444909052925 (2010).

53 Agirre, J. et al. Privateer: software for the conformational validation of carbohydrate structures. Nat Struct Mol Biol 22, 833–834, doi:10.1038/nsmb.3115 (2015).

