## Supplementary Materials for "Structures of the active HER2/HER3 receptor complex reveal dynamics at the dimerization interface induced by binding of a single ligand"

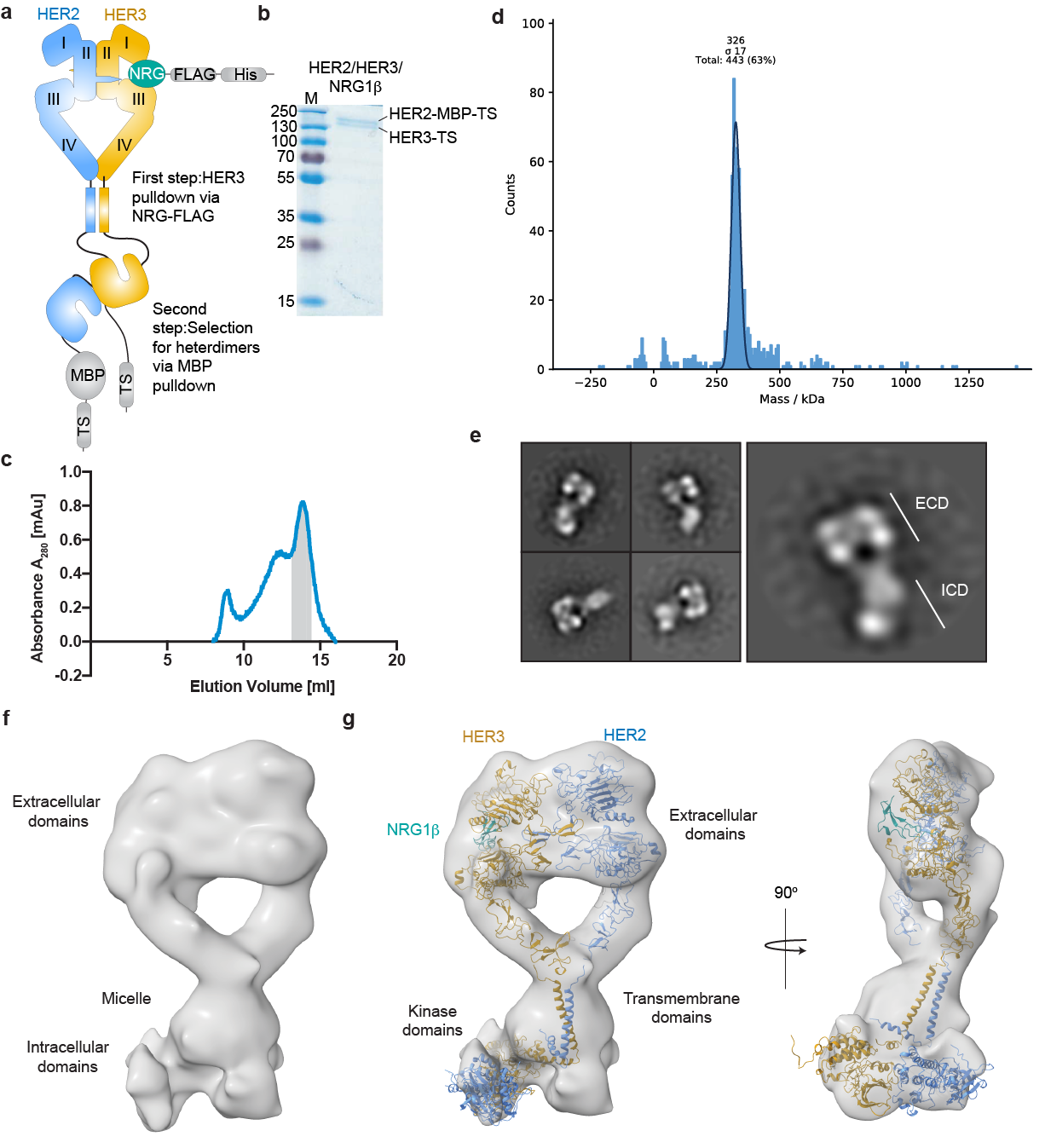

**Extended Data Fig. 1 |** **Purification, characterization, and near full-length reconstruction of the HER2/HER3/NRG1β heterocomplex**. **a,** Schematic summary of the HER2/HER3/NRG1β complex purification strategy. **b,** Coomassie-stained SDS-PAGE gel analysis of the HER2/HER3/NRG1β complex after purification showing bands corresponding to the HER2 and HER3 receptors. **c,** Representative size exclusion chromatography profile of the HER2/HER3/NRG1β complex resolved on a Superose6 10/300 Increase column (GE Healthcare). The major peak at ~14 ml elution volume is marked grey and corresponds to the fractions used for structural studies. We routinely obtained a 1-2 mAu peaks for the complex preparations from 120 ml of mammalian culture which was sufficient for the structural studies. **d,** Mass photometry of the peak sample indicates that the majority of particles have an average mass of ~326 kDa with a standard deviation of 17 kDa. The mass is consistent with that of the HER2/HER3/NRG1β complex (predicted ~280 kDa without accounting for micelle mass, ~340 kDa assuming a ~60 kDa DDM micelle mass). **e,** Representative negative stain electron microscopy 2D class averages of sample from HER2/HER3/NRG1β complex fractions (ECDs (extracellular domains), ICDs (intracellular domains)). **f,** Near full-length reconstruction of HER2-S310F/HER3/NRG1β after particle recentering with center of mass at the micelle in RELION from a ~45,000 particle stack. Rod-shaped density consistent with the asymmetric kinase domain dimer is visible below the micelle. **g,** The reconstruction accommodates models of the HER2-S310F/HER3/NRG1β extracellular domains from this study, two helical transmembrane domains and juxtamembrane-A (JM-A) segments (PDB: 2N2A), and kinases arranged in the asymmetric dimer (homology model generated from PDB: 3KEX and 3PP0). HER2 is colored in blue, HER3 in golden yellow, and NRG1β in teal.

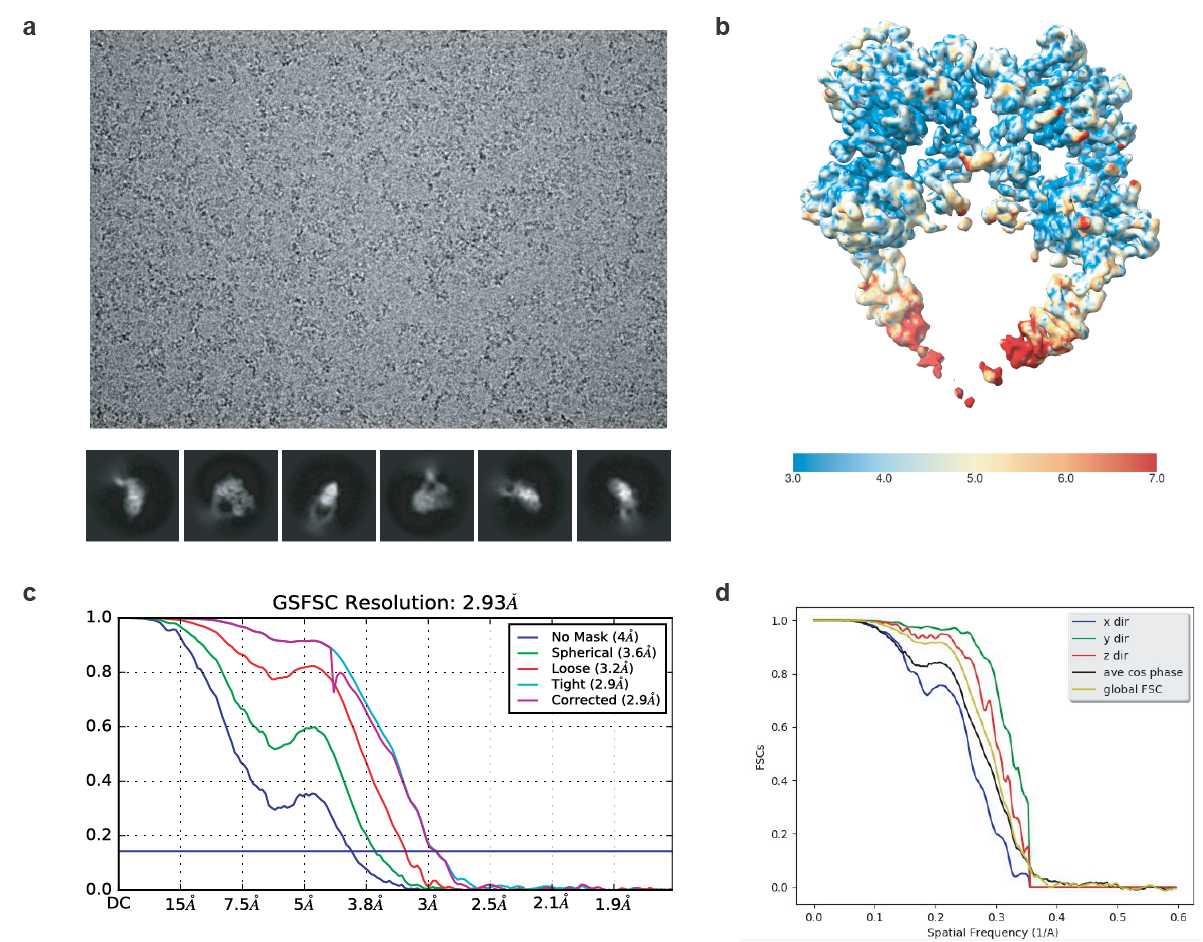

**Extended Data Fig. 2 |** **Resolution estimation and map quality of the HER2/HER3/NRG1β heterocomplex.** **a,** Top, example micrograph of HER2/HER3/NRG1β sample on graphene oxide grids. Bottom, example 2D cryo-EM class averages. **b,** Local resolution estimation from ResMap. **c,** Gold Standard Fourier Shell Correlation (GSFSC) of the final map used for model building from CryoSPARC2 with a reported resolution of 2.93 Å. **d,** Directional FSCs calculated by 3DFSC server. Map sphericity was calculated to be 0.927.

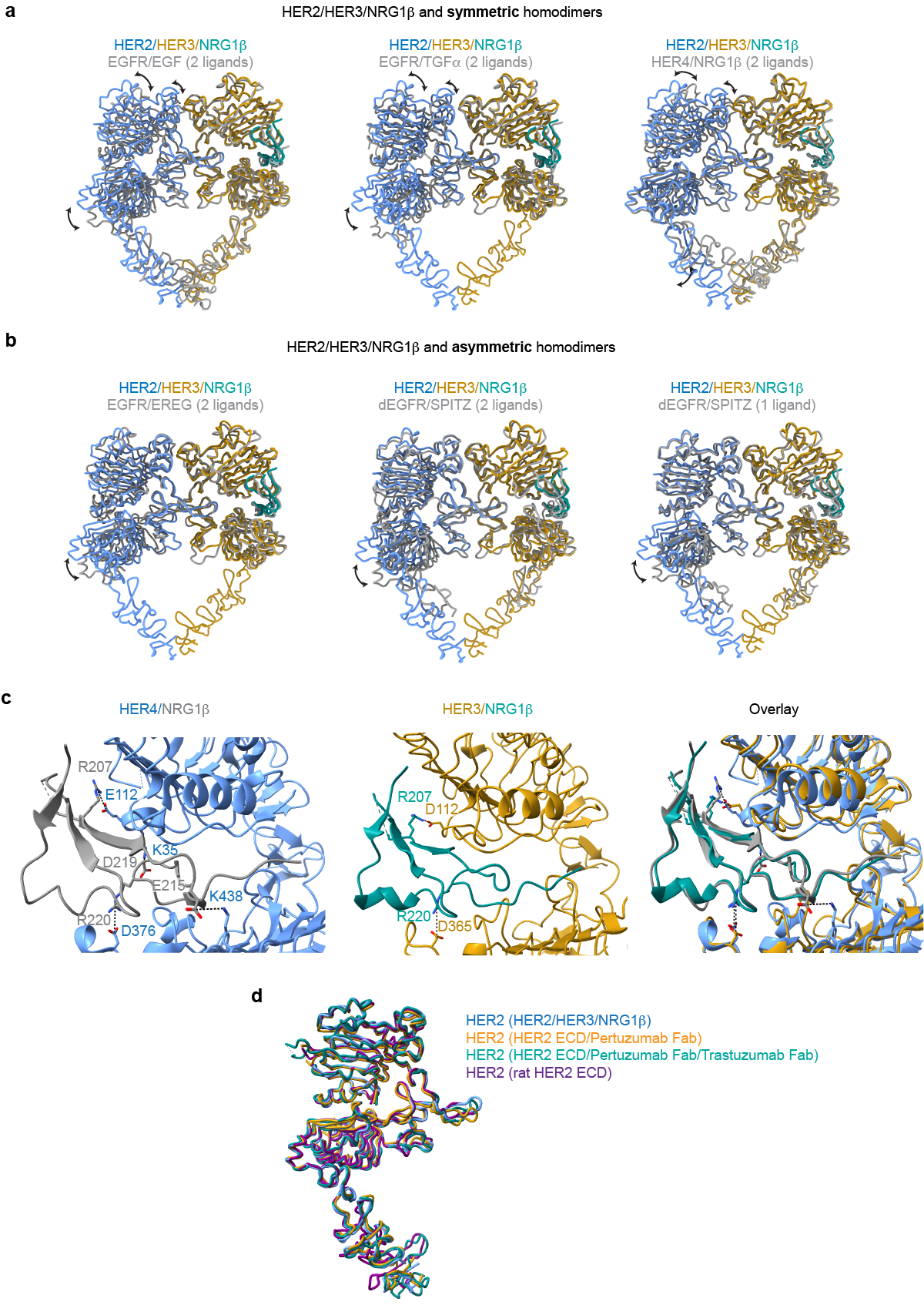

**Extended Data Fig. 3 |** **Structural comparison of the HER2/HER3/NRG1β heterocomplex with crystal structures of previously reported HER receptor structures.** **a,** Overlay of the HER2/HER3/NRG1β heterocomplexes with symmetric structures of EGFR/EGF (PDB: 3NJP), EGFR/TGFα (PDB: 1MOX) and HER4/NRG1β (PDB: 3U7U). **b**, Overlay of the HER2/HER3/NRG1β heterocomplexes with asymmetric structures of EGFR/EREG (PDB: 5WB7), doubly liganded dEGFR/SPITZ (PDB: 3LTF) and singly liganded dEGFR/SPITZ (PDB: 3LTG). All structures were aligned on HER3. Differences between the heterodimer and the homodimers are primarily appreciated in overlays on the HER2 monomer. The heterodimer more closely resembles asymmetric homodimers than symmetric homodimers but reflects a unique conformation that is not seen in previous structures. **c,** Left, HER4 bound to NRG1β (PDB: 3U7U) with salt-bridge interactions highlighted. Middle, HER3 bound to NRG1β with salt bridge interactions highlighted. Right, overlay between the two structures shows that the overall orientation of the ligand and some salt-bridge interactions are shared between HER3 and HER4, but overall HER3 forms fewer salt bridges with NRG1β than HER4. **d,** Structure of HER2 in the heterocomplex is overlayed with the crystal structure of the HER2 extracellular domain bound to trastuzumab Fab (PDB: 1N8Z), the cryo-EM structure of the HER2 extracellular domain bound to pertuzumab and trastuzumab Fabs (PDB: 6OGE) and the crystal structure of the rat HER2 extracellular domain (PDB: 1N8Y). The structures are almost identical with root mean squared deviations (RMSDs) at or below 1 Å. Minor conformational changes are observed in the dimerization arm and domain IV.

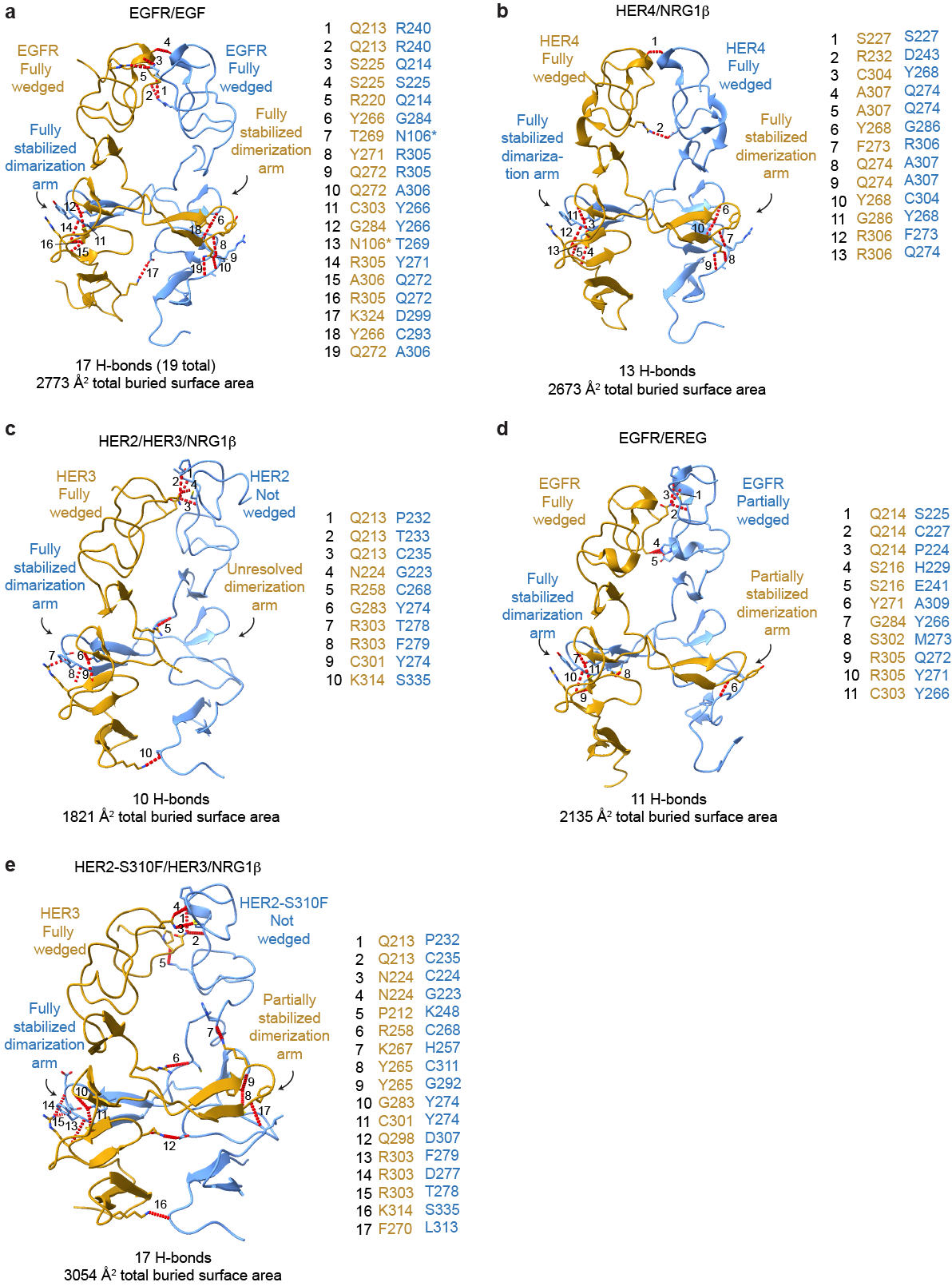

**Extended Data Fig. 4 |** **Comparison of the domain II dimerization interface between HER2/HER3/NRG1β complex domain with crystal structures of previously reported HER receptor homodimers. a-e,** The domain II interfaces of select HER receptor dimers are shown with the number of hydrogen bonds and the total buried surface area between domains I-III indicated below. Domain IV was excluded from this analysis because it is not resolved in all structures. Hydrogen bonds are shown as red dotted lines, highlighting more substantial interfaces for symmetric homodimers (EGFR/EGF (PDB: 3NJP), HER4/NRG1β (PDB: 3U7U) than asymmetric dimers (HER2/HER3/NRG1β, EGFR/EREG (PDB: 5WB7)) with the exception of the mutant HER2-S310F/HER3/NRG1β heterocomplex in which the mutation stabilizes the domain II interface. All interface hydrogen bonds are formed within domain II, except for an additional hydrogen bond in EGFR with domain III which is not shown here and marked (*) in table.

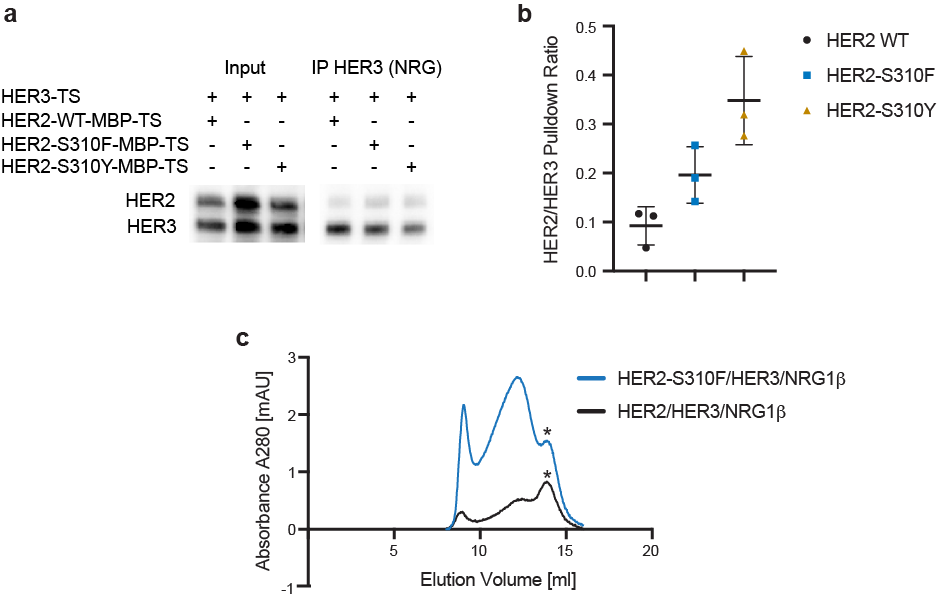

**Extended Data Fig. 5 |** **Increase in yields of the purified HER2/HER3/NRG1β complex by the oncogenic HER2-S310F/Y mutations.** **a,** Representative Western blot of ligand pulldowns of HER3 co-expressed with either HER2 WT, HER2-S310F or HER2 S310Y. **b**, Densitometry analysis of blots shown in a from three biological replicates for each condition. Error bars represent standard deviation. **c**, Overlay of representative size exclusion chromatogram profiles from a Superose6 10/300 Increase column (GE Healthcare) of WT and oncogenic HER2-S310F heterocomplexes. Heterodimer peaks are denoted by asterisks.

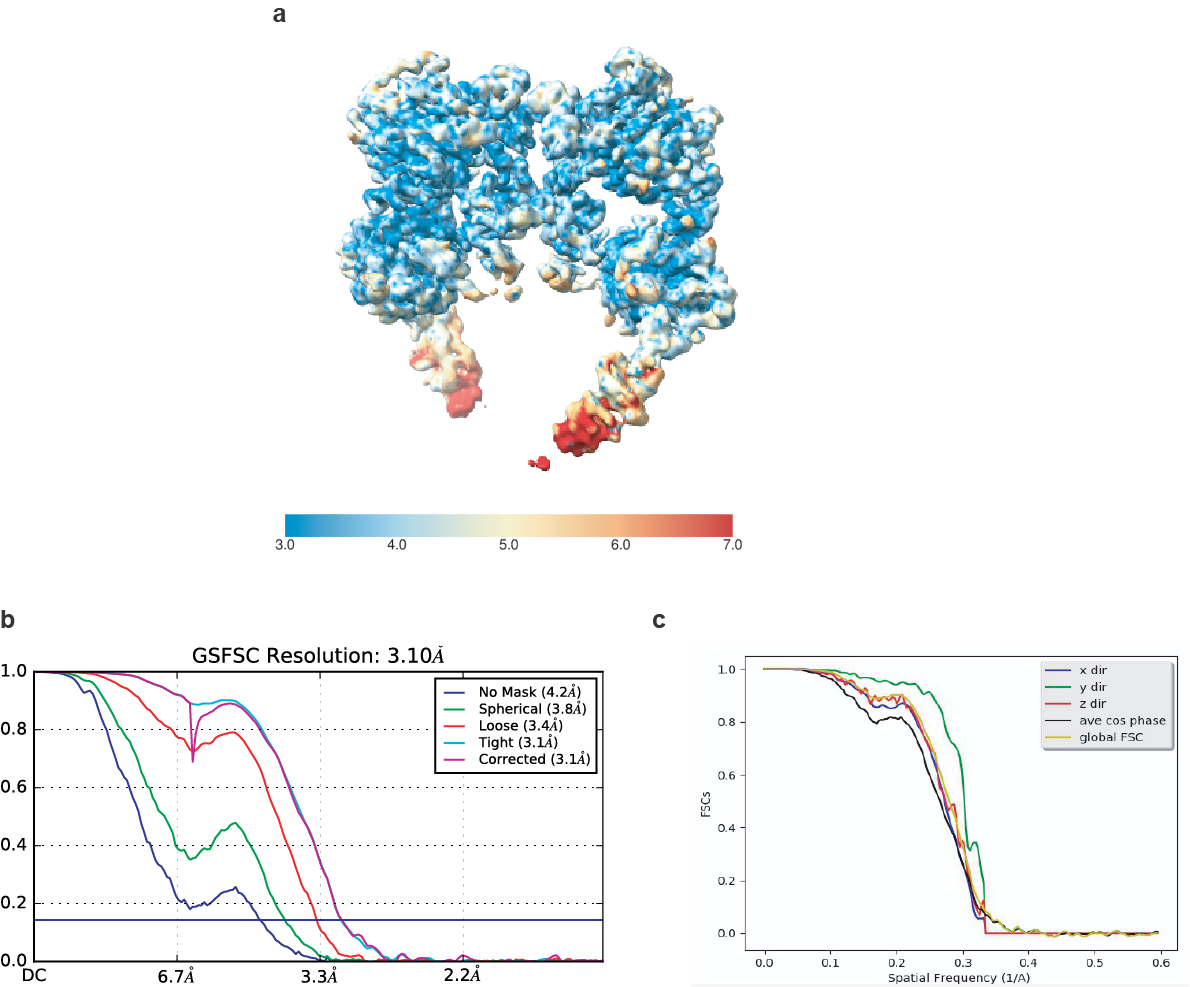

**Extended Data Fig. 6 |** **Resolution estimation and map quality of the HER2-S310F/HER3/NRG1β heterocomplex**. **a,** Cryo-EM map colored by local resolution as estimated from ResMap. **b,** Gold Standard Fourier Shell Correlation (GSFSC) of the final map used for model building from CryoSPARC2 with a reported resolution of 3.10 Å. **c,** Directional FSCs calculated by 3DFSC server. Map sphericity was calculated to be 0.949.

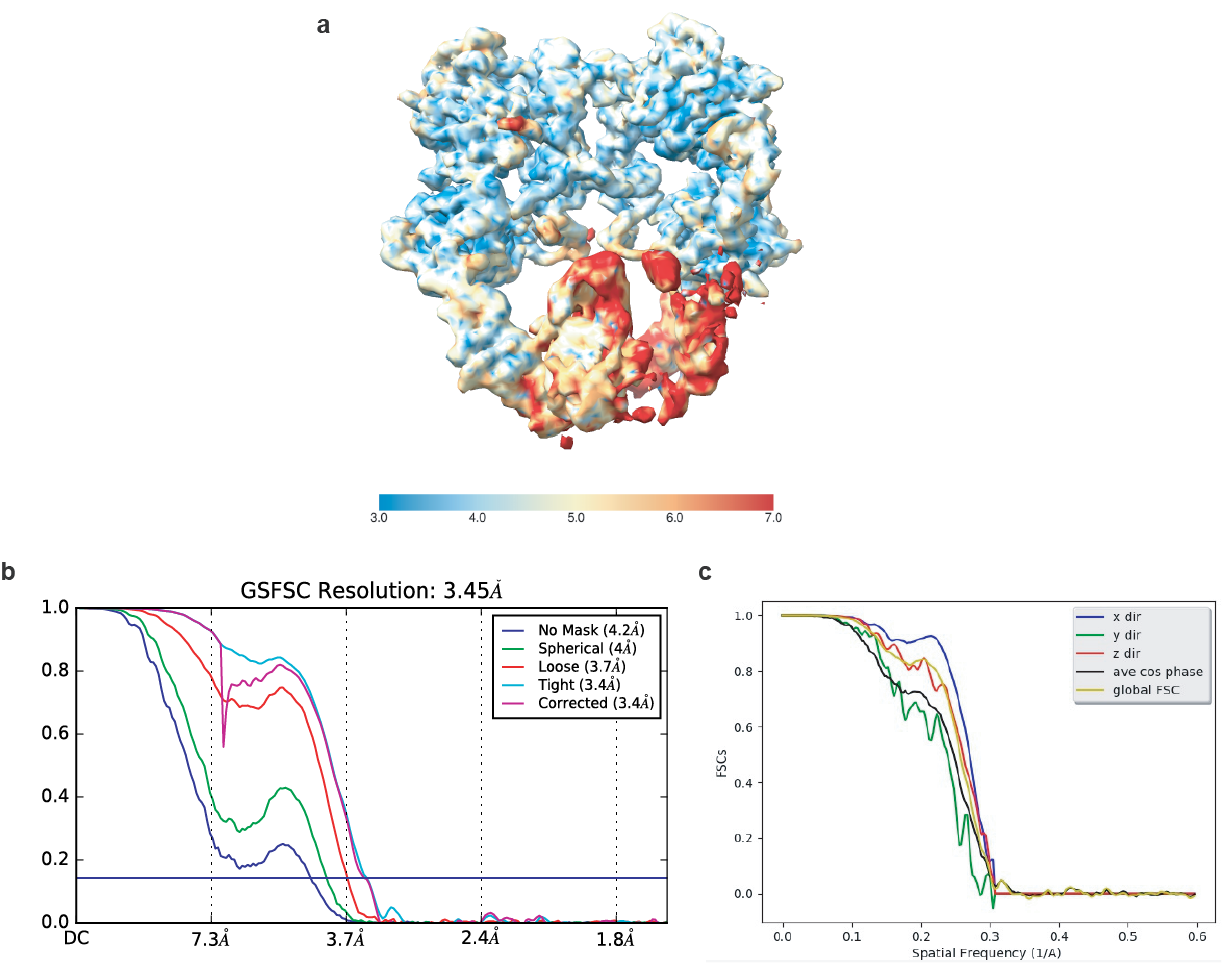

**Extended Data Fig. 7 |** **Resolution estimation and map quality of the HER2-S310F/HER3/NRG1β/Trastuzumab Fab complex**. **a,** Cryo-EM map colored by local resolution as estimated from ResMap. **b,** Gold Standard Fourier Shell Correlation (GSFSC) of the final map used for model building from CryoSPARC2 with a reported resolution of 3.45 Å. **c,** Directional FSCs calculated by 3DFSC server. Map sphericity was calculated to be 0.941.

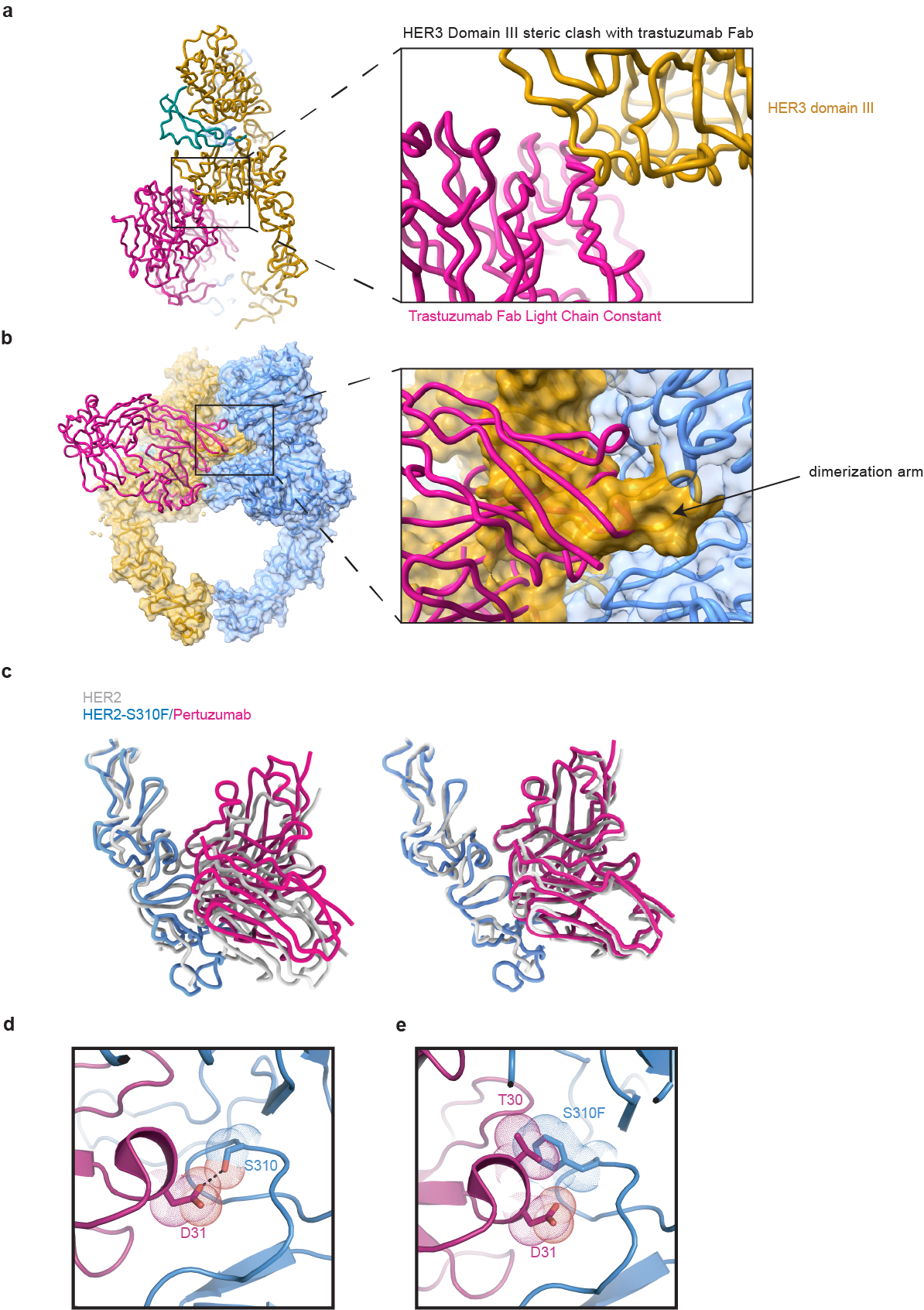

**Extended Data Fig. 8 | Docking of trastuzumab and pertuzumab Fabs in the HER2/HER3/NRG1β complex reveal steric clashes a,** Docking trastuzumab Fab (PDB: 6OGE) reveals a steric clash between the light chain constant domain with HER3 domain III. **b,** Docking pertuzumab Fab into HER2-S310F/HER3/NRG1β (PDB:6OGE) reveals a steric clash between the Fab variable domains, the HER3 dimerization arm, and HER3 domain II. **c,** Left**,** the trastuzumab Fab variable domains engaged with HER2 alone are shown from a previously published crystal structure (PDB: 1N8Z, grey) superimposed with the trastuzumab Fab engaged with the HER2/HER3/NRG1β heterocomplex (HER2 domain IV in blue, trastuzumab Fab variable domains in magenta). The two structures were aligned on HER2 domains I-III. Right, the domain IV + Fab variable domain pieces were then aligned to each other (RMSD: 1.569 Å) demonstrating a consistent epitope. Trastuzumab therefore avoids a steric clash with HER3 by inducing a rigid body rotation of HER2 domain IV relative to HER3. **d,** D31 of the pertuzumab variable light chain forms a polar contact with HER2 S310. **e,** HER2-S310F removes the polar contact with D31 and introduces a steric clash with T30 of the pertuzumab variable light chain. The predicted net effect of the oncogenic HER2-S310F mutation would be a decrease in pertuzumab affinity for HER2.

**Table 1 | Data collection and model statistics**

|  | **HER2/HER3/**  **NRG1β** | **HER2-S310F/HER3/**  **NRG1β** | **HER2-S310F/HER3/NRG1β + Trastuzumab Fab** |
| --- | --- | --- | --- |
| **Data collection** |  |  |  |
| Microscope | Titan Krios | Titan Krios | Titan Krios |
| Voltage (keV) | 300 | 300 | 300 |
| Nominal Mag | 105000x | 105000x | 105000x |
| Exposure navigation | Stage position/beam  and image shift | Stage position/beam  and image shift | Stage position/beam  and image shift |
| Cumulative dose (e^-^/Å^2^) | 67 | 67 | 66 |
| Requested defocus range (um) | 0.9-2.0 | 0.9-2.0 | 0.9-2.0 |
| Detector | Gatan K3 | Gatan K3 | Gatan K3 |
| Detector Operation Mode | CDS | CDS | CDS |
| Pixel size (physical pixel, Å) | 0.835 | 0.835 | 0.834 |
| Dose rate (e⁻/physical pixel/sec) | 8 | 8 | 8 |
| Total exposure time (sec) | 5.9 | 5.9 | 6 |
| Exposure per frame (sec) | 0.05 | 0.05 | 0.05 |
| Micrographs collected | 5228 | 4927 | 7566 |
| **Reconstruction** | EMD-23916 | EMD-23917 | EMD-23918 |
| Initial particles used | 800000 | 650000 | 1500000 |
| Particles selected after 2D classification | 200000 | 160000 | 330000 |
| Particles used in final 3D reconstruction | 123173 | 99755 | 243376 |
| Symmetry Imposed | C1 | C1 | C1 |
| Map Res (Å), masked/unmasked | 2.9/4.0 | 3.1/4.2 | 3.4/4.2 |
| FSC Threshold | 0.143 | 0.143 | 0.143 |
| Resolution range (local), Å | 3-7 | 3-7 | 3-9 |
| Final B-factor applied | -94.3 | -89.8 | -100.3 |
| **Model Refinement** | PDB-7MN5 | PDB-7MN6 | PDB-7MN8 |
| Initial Model (PDB) | 1N8Z, 1M6B, 3U7U | 1N8Z, 1M6B, 3U7U | 1N8Z, 1M6B, 3U7U |
| Protein residues (atoms) | 1211 (18325) | 1225 (18579) | 1659 (25080) |
| Ligands (atoms) | 13 (250) | 13 (250) | 13 (250) |
| Map Correlation Coefficient (masked) | 0.80 | 0.77 | 0.82 |
| RMSD, Bond Lengths (Å) | 0.010 | 0.013 | 0.013 |
| RMSD, Bond Angles ( ͦ ) | 0.963 | 1.136 | 1.056 |
| Ramachandran Outliers (%) | 0.42 | 0.28 | 0.30 |
| Ramachandran Allowed (%) | 2.93 | 2.39 | 1.77 |
| Ramachandran Favored (%) | 96.65 | 97.52 | 97.93 |
| MolProbity score | 1.15 | 1.03 | 0.89 |
| Clashscore (all atoms) | 1.78 | 1.75 | 1.38 |
| Rotamer outliers (%) | 0.10 | 0 | 0 |
